# Using pangenome variation graphs to improve mutation detection in a large DNA virus

**DOI:** 10.1101/2025.11.26.690900

**Authors:** Tim Downing, Chandana Tennakoon, Lidia Lasecka-Dykes, Caroline Wright

## Abstract

Accurately quantifying viral genetic diversity is essential for understanding pathogen evolution, transmission, and emergence. However, standard approaches that map sequencing reads to a single linear reference genome introduce substantial reference bias, particularly for samples that are divergent, recombinant, or belong to rare lineages. Pangenome variation graphs (PVGs) mitigate this issue by representing multiple genomes within a unified graph structure, enabling read mapping across all observed and potential haplotypes. Despite this, PVGs have rarely been applied to viruses. Here, we address this gap by constructing and evaluating the first PVG for lumpy skin disease virus (LSDV), an emerging poxvirus of global importance. We generated PVGs of different sizes and mapped Illumina datasets using Giraffe, benchmarking performance against linear reference mapping with Minimap2. A minimal three-sample PVG containing one representative from each major lineage recovered 97% of known LSDV nucleotide diversity while reducing PVG size by >95% relative to a 121-sample PVG. PVG-based mapping detected more SNPs than linear mapping, including variants supported by read evidence that was not detected when reads were mapped to a single reference genome. 27% of SNPs detected using PVG-based mapping could not be projected onto the linear reference coordinate system because they were on alternative paths absent from reference, highlighting the impacts of reference bias. Notably, these new SNPs were at genes involved in host recognition and immune evasion, and identified lineage-specific mutations that improved subclade phylogenetic structure. Our findings demonstrate that PVGs substantially enhance SNP discovery in LSDV, with direct implications for genomic surveillance, outbreak tracing, and the detection of recombinant vaccine-related lineages in LSDV and other large DNA viruses.

**Impact Statement:** Genomic surveillance of viral pathogens typically relies on mapping reads to a single reference genome, a practice that systematically misses variation in divergent or recombinant isolates. We show that pangenome variation graphs (PVGs) can reduce this limitation in DNA viruses. Using lumpy skin disease virus (LSDV) as a model system, we demonstrate that a compact PVG constructed from only three representative genomes captures most known genomic diversity and improves the detection of mutations that are difficult to identify using linear reference mapping. PVGs recover biologically meaningful variants at genes involved in immune evasion and host interaction, resolve closely related subclades, and reveal mutations that cannot be identified using a single reference genome. This work uses a population-structure-guided strategy for building reference viral PVGs that balance computational efficiency with sensitivity, offering clear benefits for genomic surveillance, outbreak reconstruction, and the detection of recombinant lineages.

**Data Summary:** All supporting code and protocols have been provided within the article or through supplementary data files. Supplementary tables are available at Figshare: https://figshare.com/s/7a879df5b5452d04585b with all code at https://github.com/downingtim/LSDV_pangenomics

## Introduction

Traditional genomic analyses rely on mapping sequence reads to a single linear reference genome. However, this approach introduces reference bias when the sequencing reads differ from the reference, causing mismatched reads to align poorly and resulting in missed or mis-called SNPs, as well as structural and copy number variants [1–4]. Such biases reduce the accuracy of downstream phylogenetic reconstruction and genotype–phenotype association [5,6]. Importantly, neither consensus nor multi-sample linear references have overcome these limitations in viral systems [7,8].

Pangenome variation graphs (PVGs), also known as pangenome graphs or sequence graphs, address this reference bias by representing multiple genomes together in a single graph structure that encodes the nucleotide-level diversity of a sample collection [1]. PVGs represent sequences as nodes connected by edges, where a path through the graph corresponds to the genome sequence of an individual sample.PVGs have proven effective in human genomics, improving variant detection and reducing reference bias, particularly at structurally variable regions [1,9]. Their key advantage is that they reveal additional variants, including biologically meaningful changes that are invisible to linear reference approaches. Importantly, PVG-based read mapping is now computationally efficient, sensitive and accurate [1,9–13].

Viruses exhibit diverse genome structures and evolutionary behaviours, broadly grouped into the seven Baltimore classes [14]. RNA viruses (including segmented ones) typically show high mutation rates [15] and extensive recombination [16] or reassortment [17], generating deep within[species diversity that is difficult to represent comprehensively in a single reference. In contrast, large dsDNA viruses such as poxviruses evolve more slowly but can accumulate substantial structural changes, gene duplications, and recombination-derived mosaics [18]. These class[specific processes influence how species[level diversity is distributed, and in many cases make graph[based approaches such as PVGs better suited than linear references for capturing both shared and lineage[specific variation.

PVGs have seen limited use in viral genomics so far [19]. One notable exception is the PVG constructed for the circular 3.2 Kb hepatitis B virus, which successfully mapped more than 8% of reads that failed to align to linear references, particularly in highly diverse samples [10]. Viruses are relatively short, which reduces computational overheads, and recent PVG methods now support genomes of any length [20]. PVGs are well suited to viral datasets because they capture variation associated with phenotype, recombination, and rare divergent lineages, features that often pose major challenges for linear reference approaches. These advantages are especially valuable for tracking recombinant vaccine-derived lineages in livestock viruses.

Here, we demonstrate how PVG-based read mapping can improve inferences compared with mapping to a linear reference, using Lumpy skin disease virus (LSDV) as an exemplar. LSDV is one of three members of the Capripoxvirus genus within the Poxviridae. It causes severe economic losses by infecting cattle, resulting in reduced milk production, hide damage, and mortality [21–23]. LSDV infection is a notifiable disease by the World Organization for Animal Health (WOAH). Over the past two decades, LSDV has spread from sub-Saharan Africa into the Middle East, Asia, and more recently Southeast Asia and China [24–27]. LSDV is an emerging virus and accurate measurement of its genetic diversity is essential for linking evolution to geographic spread and drivers of socio-economic impact.

The LSDV genome is ~150 kb in length, AT-rich (75%), and typically encodes 156 genes [28]. It has three major recognised phylogenetic lineages: vaccine-related Clade 1.1, wild-type Clade 1.2, and recombinant Clade 2 [29–30]. Although LSDV evolves more slowly than RNA viruses, extensive recombination between wild-type and vaccine strains has generated mosaic haplotypes with altered phenotypes and possible vaccine-escape properties [31–33]. These mosaics cannot be fully represented by a single linear reference genome [34–35]. Furthermore, LSDV genomic diversity is elevated at genes involved in host recognition and immune evasion, which are typically located near the genome ends and require accurate variant detection. Such variation is critical for outbreak source tracing, monitoring recombinant lineages, and understanding mutations that influence cattle immune responses.

Here, we tested whether a PVG-based read mapping strategy improves variant detection compared with linear reference mapping, using LSDV as a model. We constructed PVGs from all available LSDV genome assemblies using existing tools [20] and mapped both real and simulated Illumina libraries to multiple PVG versions using Giraffe [12] and, for comparison, to a linear reference using Minimap2 [36]. We therefore tested whether a small set of representative genomes chosen from the major phylogenetic lineages could produce a compact PVG that retains most detectable diversity while remaining computationally efficient. This approach was computationally scalable, enabled rapid PVG-based analyses and can be generalised to other viral systems. This revealed additional mutations and provided improved insights into LSDV genetic diversity from evolutionary, phylogenetic and functional perspectives.

## Methods

### Genome assembly collection, alignment, phylogenetic analysis, quality control and annotation

All available complete LSDV genomes (n=128) were downloaded from the NCBI nucleotide database [37] (15^th^ November 2023), from which the first and last 300 bp of the genome were masked. This initial dataset was aligned with Mafft v7.453 [38] using automatic optimisation and default parameters (Figure 1).

**Figure 1.**
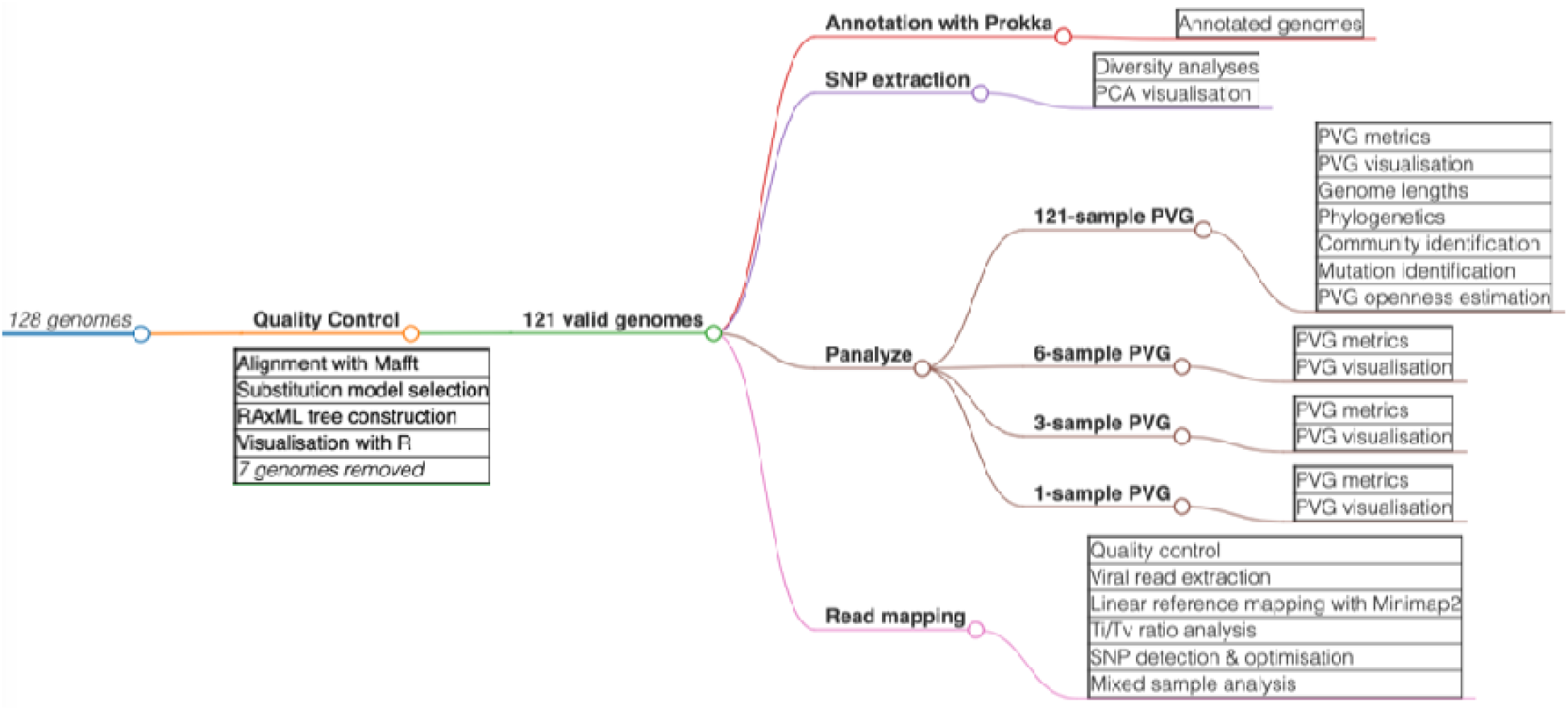
Methods summary showing the flow of data and analyses. After quality control of the initial dataset of 128 genomes, 121 validated genomes were subject to annotation, recombination detection, diversity analyses, PCA, phylogeny construction, PVG creation from 121-, 6-, 3- or 1-sample sample/s, and association PVG investigation (coordinate comparison, annotation, visualisation, community detection, and comparisons across genomic regions).

Evolutionary relationships among these genomes were reconstructed using RAxML-NG (Randomised Axelerated Maximum Likelihood) v1.2.0 [39] with a GTR (general time reversible) model and gamma substitution rate heterogeneity selected by modeltest-ng [40]. These phylogenies were midpoint-rooted and visualised using the following R v4.3.2 [41] packages: ape v5.7-1 [42], ggtree v3.8.2 [43], phangorn v2.11.1 [44], Rcpp v1.0.11 [45], RcppArmadillo v0.12.6.6.0 [46], phytools v2.0-3 [47] and treeio v1.24.3 [48].

Seven genomes were not included because their assemblies were reverse complemented [49], suggesting the genome assembly quality was low. Of these seven genomes, OK422493 (Ranchi 1 P30, 31/12/2019, India), OK422492 (Ranchi 1 P10, 31/12/2019, India) and ON400507 (208 PVNRTVU, 2020, India) were nearly genetically identical and were previously classified as LSDV Clade 1.2.2 (Breman et al 2023), and OM373209 (BH3 CHN 20, China) was previously classified as recombinant LSDV Clade 2.5.1 (Breman et al 2023). The three others were OM984486, OQ606832 (NIAB, PVNRTVU, India, 2022) and OQ427097 (West Bengal, 2022, India).

This resulted in a high-quality dataset of 121 genomes, which were annotated using Prokka v1.14.6 [50] using the LSDV isolate KX894508 (155920/2012) annotation that had 163 CDS regions (Figure 1). The median genome length was 150,575 bp: MW631933 (Morocco, 14/07/2017) was the sole sample with a length <150 Kb. The relative difference in genome coordinates across the 121 samples was small: the median coordinate difference compared to KX894508 was 109 bp (Figure S1). MW631933 was the only sample with coordinates >360 bp different to the others.

### Genome partitioning, SNP analysis, and phylogenetic resolution

Alignment and phylogenetic resolution of the 121 genomes were implemented as above (Figure 1). This was also applied for the core, 5’ accessory and 3’ accessory genomic regions. The core region was defined as a conserved 93 Kb region between bases 13,851 (the start of gene LD020, encoding a ribonucleotide reductase small subunit) and 106,910 (before gene LD116, encoding an RNA polymerase subunit), inclusive, as per previous work [29]. The 5’ accessory genome extended to base 13,850 (13,850 bp in total). The 3’ accessory genome was designated from base 106,911 onwards, with the core region at 13,851-106,910 bp.

The SNPs were extracted per sample as VCFs using SNP-sites v2.5.1 [51]. Comparisons and analyses of sequence diversity across groups and between sample pairs was implemented with VCF-kit v0.2.9 [52] and vcflib [53]. The core region had five triallelic sites, the 5’ end had one such site and the 3’ end had four triallelic sites. We performed principal component analysis (PCA) using R package adegenet v2.1.10 [54] based on genome-wide SNP data (Figure 1).

### Pangenome variation graph creation

We used Panalyze [20] to implement PVG construction and analysis. We used it to make four PVGs (Figure 1): one with all 121 assembled genomes, another with three representing the three major clades, one with six (spread geographically and over time), and one with a single sample (KX894508). In make the latter one-sample PVG, construction requires two or more sequences, so the KX894508 genome was duplicated and modified at a single base to create a minimal PVG. KX894508 was the linear reference genome. Panalyze created PVGs in Graphical Fragment Assembly (GFA) format using the pangenome graph builder (PGGB) v0.5.4 pipeline [55] via all-to-all alignment with wfmash v0.7.0 [56]. Multiqc v1.14 [57] was used to collate and assess PVG metrics. These PVGs were converted with “convert” from variation graph (VG) v1.43.0 [9], and indexed with VG autoindex. The corresponding genome files were indexed with SAMtools [58].

An advantage of using graphs is the ability to perform computationally scalable indexing through bidirectional graph Burrows Wheeler Transform (GBWT) indices, which store haplotype paths as nodes [13,59]. Consequently, we created GBWT-indexed versions of our PVGs with VG gbwt using the default minimiser length of 29 and window size of 11 for the Giraffe minimiser index. The GBWT indices were created using the greedy path-cover algorithm across all PVG paths and were converted with VG. Then VG snarls was used to create a snarls file. This was used to create a distance index of the PVG’s snarl decomposition so the minimum distances between nodes in the PVG were calculated. Next, VG minimizer used these distances to identify each k-mer’s position within the PVG for each haplotype, along with their corresponding PVG distances. VCFs were obtained from VG deconstruct using the GBWT-indexed graphs.

### Pangenome variation graph manipulation, visualisation and annotation

Panalyze analysed these PVGs using ODGI (optimized dynamic genome/graph implementation) v0.8.3 [60], including quantification of SNPs, indels and compound mutations using VG deconstruct. Panalyze used gfautil v0.3.2 [61] to get mutation rates and coordinates for genomes at each path and site. Subgraphs were selected using gfatools v0.4 [62], evaluated with gfastats v1.3.6 [63], analysed with gfastar v0.1 [63], and visualised with waragraph [64] and VG. Presence-absence variants (PAVs) were identified and visualised using ODGI via Panalyze and the R package ggplot2 v3.4.4 [65].

The PVGs were annotated using Prokka [50] and were indexed with SAMtools v1.19 [58]. Using the reference genome annotation for KX894508, Busco v5.5.0 [66] found a mean of 155.8 complete unique CDSs per genome (standard deviation 1.5). The Prokka GFFs were converted to gene transfer format (GTF) with gffread [67] and parsed. The annotation position information was merged with the ODGI rendering of the PVGs, creating one CSV annotation file per PVG, which was visualised using Bandage v0.8.1 [68].

### Community detection and visualisation

PVG communities reflected the relative similarity of genomes based on pangenome data for a specific region. Here, these were identified via Panalyze’s implementation of pairwise alignment using wfmash [56] via PGGB [55] with a k-mer of 19 bp and a window size of 67 bp, subject to thresholds of 80% sequence similarity using mash distance and ten mappings per node. Panalyze indexed the file names using fastix [69], and evaluated these communities with PGGB scripts [55]. This built a network with the contigs as nodes and mappings as weighted edges, such that the weights were proportional to the percentage identity of the mapping and its length. These networks were visualised with igraph tools [70].

### Read library sample collection and quality control

A total of 76 Illumina read libraries were downloaded from the SRA (24^th^ May 2024) and six additional unpublished datasets were added to make a total of 82. The metadata for each library was retrieved where available, including host species, year/country of isolation, sequencing platform, library source/selection and preparation approach (Table S1).

Adapter trimming, removal of reads with low base quality (BQ) scores (phred score <30), exclusion of ambiguous (N) bases, primer removal, correction of mismatched bases in overlapping regions, and trimming of poly-G tracts at 3′ ends was completed using Fastp v0.23.4 [72]. Low-quality bases with BQ <30 and reads with lengths <110 bp were removed using the FASTX-Toolkit v0.0.13 [73]. FastQC v0.12.1 [74] and MultiQC v1.14 [52] were used to assess the effectiveness of Fastp and the FASTX-Toolkit. A total of 89% of Illumina read library bases had BQ>30, similar to previous work [75]. The number of screened reads was higher for metagenomic libraries (mean 36,146k) compared to WGS (3,063k) and amplicon (678k) libraries (Table S2). The fraction of reads excluded because they were low quality or had too many Ns was similar across library types. Reads that were too short or adapter-trimmed were only found in WGS libraries. Median read duplication rates were higher for amplicon (19.0%) and WGS (18.8%) compared to metagenomic libraries (10.9%).

We estimated species-level abundances for each read in each library using Kraken2 [76] with the core nucleotide database. All libraries had reads stemming from *Bos, Cervus, Homo* and *Ovis* in addition to LSDV (online Supplementary Material 1). The relative numbers of reads allocated to each stage were assessed (Table S3).

### Pangenome variation graph read mapping, testing and comparisons

We mapped the viral reads of each library using Giraffe in VG v1.43.0 [9] to each PVG to generate Graph Alignment Map (GAM) files. Giraffe mapped each read library to the PVG containing one (named Giraffe_1 here), three (Giraffe_3), six (Giraffe_6) and all (Giraffe_all) samples. VG uses efficient PVG indexing and read mapping using a partial order alignment [2]. The PVGs were initially in XG format (topological representation) with accompanying GCSA2 indices for read mapping using VG-MAP. To enable mapping with Giraffe, the PVGs were converted from GFA to GBZ format, which includes a compressed graph and an index of all haplotypes. The GBZ file was indexed with VG, and the minimizer distances were used by VG call. VG surject was used to create Binary Alignment Map (BAM) files, which represent the reads at their reference nodes in the PVG. We obtained GAM read mapping statistics across all snarls using VG stats. VG paths and VG mod were used to extract haplotype paths from the individual graphs. Mapping with Giraffe was much better than VG-MAP [13], so the latter is not discussed further.

We used GAM files to visualise read mapping patterns of individual samples on the PVGs. Augmented graphs were then generated using VG augment, with the following thresholds: a minimum path coverage of three, a base quality (BQ) of at least 10, and a minimum mapping quality (MQ) of 10. Snarls were interpreted based on the augmented graphs using VG snarls. VG packgenerated PACK files from the augmented graphs, which are compressed coverage indices, analogous to Sequence Alignment Map (SAM) files. Each unique snarl in the PVG was used to screen for potential mutations in VCF files by VG call, which used the PACK file and augmented graph data. Mutations were detected if the sample’s paths in the PVG traversed the allele of interest. These PACK files were used as input for VG depthto calculate path depth at nodes with a coverage of six-fold or more. The compute resources and time taken for this VG-based approach were greater than that using the surjected BAM files. To identify candidate mutations from GAM files via VG, we assumed haploidy and required variants to have a MQ of 10+. This was processed by using VG augment, snarls, pack and call to generate VCF files for valid candidate mutations (labelled with “PASS”). These VCFs were used to interpret SNPs with allele frequencies between 5% and 90%. The mapped reads in the GAM files were visualised with SequenceTubeMap [77].

We used Minimap2 v2.24-r1122 [36] to map libraries to the KX894508 genome and generate SAM files, which were converted to BAM format using SAMtools v1.11 [58]. The latter was then used to sort these BAM files, including surjected BAMs, extract information on the mate coordinates, calculate insert size, resolve missing read pairs, remove duplicate reads, index them and extract read mapping statistics. Mutations detected with VG were extracted from the corresponding PACK files. BAM file metrics were obtained using SAMtools [58] flagstat, while read depth data were derived using SAMtools coverage and was parsed with R to measure read depth, BQ and MQ.

### Mutation detection and screening

We filtered for candidate mutations (SNPs and indels) based on the BAM files using BCFtools v1.20 [58] and Freebayes v1.3.7 [78]. BCFtools detects mutations based on the frequency of derived alleles in sequencing reads and has been shown to outperform alternative options like GATK in certain contexts [79]. Freebayes identifies mutations across haplotypes using a Bayesian approach [78]. Previous studies have shown that stringent thresholds are effective for viral sequence data [80], both BCFtools and Freebayes are effective for mutation detection [81], and these tools work well with PVG-based approaches [82].

Mutations were considered valid if they met the following criteria: BQ (QUAL) score of 30+, a minimum read depth of 10, at least five reads supporting the alternate allele, and whose location was not within 2.5 Kb of the genome ends, where coverage and alignment quality were more variable. These thresholds were consistent with previous studies, and testing with a lower BQ threshold of 20 impacted the number of SNPs per sample by <1%, demonstrating no impact on key comparisons. For variant calling with BCFtools, we also required that SNPs were haploid and for Freebayes, that allele frequency was >0.05. These parameters were selected and optimised by manually inspecting candidate SNPs using IGV [83]. The resulting valid VCF files from FreeBayes and BCFtools were merged and then normalised with BCFtools to represent multiallelic sites correctly. This produced fiveVCF files per sample across mapping approaches (Giraffe and Minimap2), and PVG options (one-, three-, six-, all-sample).

Transition-transversion (Ti/Tv) ratios were calculated using BCFtools for all detected mutations before and after filtering. A neutral Ti/Tv ratio=0.5 was expected for non-conserved regions because there are twice as many mutational options for transversions compared to transitions. A Ti/Tv ratio > 2 can be indicative of functionally active or constrained regions, as observed in *Drosophila* [84] and humans [85–87]. A Ti/Tv ratio > 3 has been observed in highly constrained loci [88] including inter-species comparisons of viral genomes [10]. Although reduced Ti/Tv ratios can result from mutation saturation, here these effects may be less pronounced because we examined a slow-mutating dsDNA virus. To help identify reliable SNP-calling pipelines, we required that any valid dataset to have a Ti/Tv > 2.

Comparisons between SNP datasets were implemented using the BCFtools concat. We used the FLO toolkit (https://github.com/wurmlab/flo [85], SAMtools [53] and Picard LiftoverVcf [86] to transfer coordinates between VCF files with different reference genomes. This liftover converted mutations from the three-, six- and all-sample PVGs to the KX894508 annotation, requiring a sequence identity of >98%. We identified valid mutations with mismatched reference alleles, which indicated that the KX894508 reference lacked the alternative allele. These “unlifted” mutations represented SNPs uniquely detected using PVG-based methods that were missed by traditional linear reference-based approaches. SNP data was processed with Python and visualised using R package ggplot2 v3.4.4 [60].

The functional impact of SNPs was examined by comparing the FASTA and GenBank annotations of KX894508, OQ511520 and KX764645 against the VCF files using our LSDV variant effect predictor. This detected SNPs as causing nonsynonymous, synonymous, intergenic, stop-loss or stop-gain mutations within Clades 1.1, 1.2 and 2. We also calculated the rates of nonsynonymous and synonymous variants per sample.

### Simulated data analysis

We analysed 40 simulated paired-end read libraries based on the genomes from OQ511520 (Clade 2) and KX764645 (Clade 1.1) using art_illumina v3.19.15 [71]. These were compared to KX894508 as the main reference (Clade 1.2). Each sample had 20 libraries with read depths incremented by 10, starting at 10 and going up to 200 inclusive. These 40 libraries went through the same quality control, read mapping and SNP detection as above. This used the SNPs derived from genome alignment of OQ511520 versus KX894508 (1,130 SNPs), KX764645 versus KX894508 (1,911 SNPs), and KX764645 versus OQ511520 as the truth sets to measure the numbers of SNPs found per method differentiating OQ511520, KX764645 and KX894508. Metrics related to true positives, false positives, true negatives, false negatives, precision, recall and F1 scores were extracted from this data.

### Co-phylogenetic analysis of the effects of different mapping approaches on phylogenetic inference

We examined the phylogenetic and co-phylogenetic effects of SNP detection across different mapping strategies, directly comparing linear reference and PVG-based approaches. This used the merged SNPs from Giraffe_1 and Giraffe_3 (called Giraffe_1&3), and separately the merged SNPs from Giraffe_1 and Giraffe_6 (called Giraffe_1&6). To do this, we created a concatenated SNP alignment containing the valid SNPs for 78 samples across the six genomes in the six-sample PVG (total lengths 903.7 Kb). Four divergent samples (SRR8442780, SRR19090746, SRR19090747, SRR19090748) were excluded because they were partially non-LSDV. Maximum likelihood phylogenies were inferred using RAxML-NG v1.2.0 [39] using a GTR+G4 model. The trees were midpointed-rooted and visualised in R. We used phytools v2.0-3 [47] for cophylogenetic comparisons. The clades were annotated based on published data [29]. These alignments had comparable numbers of parsimony-informative sites for mapping to the linear reference with Minimap2 (1,181 sites), or with Giraffe to the one- or three-sample PVGs (1,041), or with Giraffe to the one- or six-sample PVGs (1,003), respectively.

## Results

### Pangenome variation graph analysis of LSDV genomes

We constructed a PVG of 121 LSDV genomes (called the all-sample PVG) with Panalyze [20]. This PVG revealed varied genetic diversity across the genome with double the SNP density at the 5’ and 3’ ends compared to the core region (Figure 2). This was supported by a higher Ti/Tv ratio at the core (Table 1). Additionally, we found that the allele frequency spectrum had an exponential decay with increasing sample count (Figure S2), indicating that most SNPs were rare and lineage-specific across the three major clades, with only a minority reaching frequencies > 50%.

**Figure 2.**
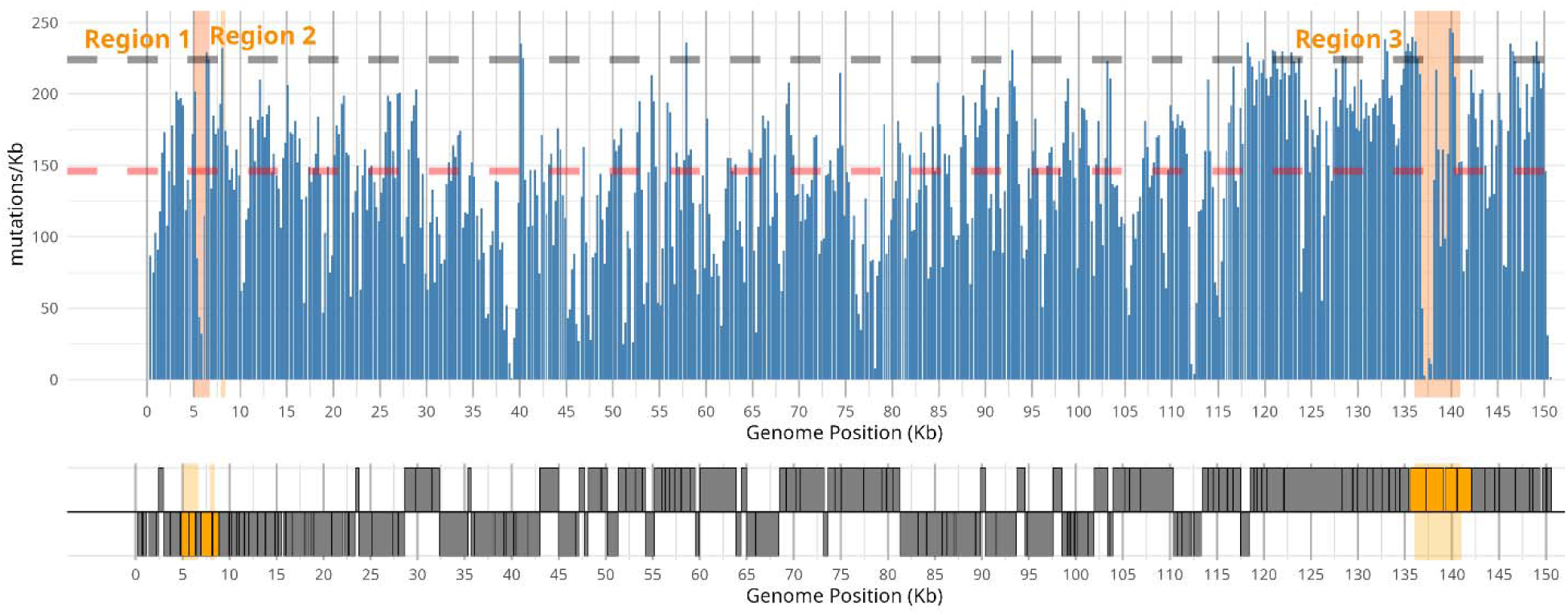
PVG-wide diversity (SNPs/Kb) in LSDV using 500 bp sliding windows (blue bars). The KX894508 genome annotation is represented below as grey boxes that are read in the forward (lower strip) or reverse (upper strip) directions. The red dashed line indicates the median number of SNPs/Kb. The gray dashed line denotes the 95% percentile. Three regions of interest and associated CDSs are in orange: (Region1) 5,200-6,500 bp, (Region2) 8,055-8,231 bp and (Region3) 136,200-140,400 bp.

**Table 1.** Genetic diversity across the whole genome, 5’ end, core region, and 3’ end. Values include the transition/transversion ratio (Ti/Tv), Watterson’s theta per Kb (Theta/Kb), and the mean number of pairwise differences per Kb (Pi/Kb) across samples.

| Region | Length (bp) | SNPs | Theta/Kb | Pi/Kb | SNPs/Kb | Ti/Tv |
| --- | --- | --- | --- | --- | --- | --- |
| 5' accessory | 13,610 | 325 | 4.1 | 6.8 | 23.9 | 2.40 |
| core region | 93,059 | 1,076 | 2.3 | 3.8 | 11.6 | 3.49 |
| 3' accessory | 43,644 | 1,107 | 4.7 | 7.6 | 25.4 | 2.37 |
| total | 150,313 | 2,436 | 3.3 | 5.4 | 16.2 | 2.89 |

### Subset-derived pangenome graphs capture broad LSDV diversity

Phylogenetic reconstruction of the PVG’s SNPs recapitulated the known population structure [32] (Supplementary Text 2), with three major clades: vaccine-related Clade 1.1, wild-type Clade 1.2 and recombinant Clade 2, along with several rarer lineages (Figure S3). We hypothesised that LSDV’s simple population structure would allow building a PVG based on a subset of samples to allow quicker computation and interpretation without affecting mutation detection. To explore this, we created PVGs from three and six genetically representative samples (referred to as the three- and six-sample PVGs) (Table 2). The three-sample PVG had a representative from each of the main clades: KX764645 from vaccine-related Clade 1.1, KX894508 from wild-type Clade 1.2, and OQ511520 from recombinant Clade 2. These samples were chosen because each had limited unique variation, was closely related to other members of their clade, and KX894508. Sequences of these isolates were also used as linear reference. Pairwise comparisons showed 1,911 SNPs between KX764645 and KX894508, 844 between KX764645 and OQ511520, and 1,130 between OQ511520 and KX894508. These differentiation rates were consistent with the wider dataset, for which the median pairwise SNP distance was 1,858 (standard deviation 1,080). For the six-sample PVG, more than one sample per clade was used to account for assembly artefacts, differences in genome completeness, and subclade-specific diversity. In Clade 2, MW732649 and OQ511520 differed by 1,272 SNPs: MW732649 had novel variants absent from the other samples. In Clade 1.2, OK318001 differed from KX894508 by 706 SNPs and from KX683219 by 907 SNPs, and KX894508 and KX683219 differed by 611 SNPs.

**Table 2.** GenBank accession and associated sample details for genomes included in the three- and six-sample PVGs.

| Accession | Sample name | Subclade | Country | Year | PVGs |
| --- | --- | --- | --- | --- | --- |
| KX894508 | 155920/2012 | 1.2.1 | Israel | 2012 | 3 & 6 |
| OQ511520 | Thailand/Trang/2022 | 2 | Thailand | 2022 | 3 & 6 |
| KX764645 | Neethling-LSD_vaccine-OBP | 1.1 | South Africa | NA | 3 & 6 |
| OK318001 | V281 | 1.2.3 | Nigeria | 2018 | 6 |
| KX683219 | KSGP_0240 | 1.2.2 | Kenya | 1974 | 6 |
| MW732649 | HongKong/2020 | 2 | Hong Kong | 2020 | 6 |

The all-sample PVG was 155,008 bp (151,506 bp excluding non-ACGT, ‘N’ bases), of which 147,614 bp were shared, indicating that 97% of the LSDV genome was conserved. By comparison, the six-sample PVG was 151,246 bp (150,643 bp without Ns), indicating that only a small amount (863 bp, 0.6%) of additional information came from the extra 115 genomes. Complexity was substantially lower in the six-sample PVG: the sum of sequence lengths was 18,213 Kb in the all-sample PVG versus 904 Kb in the six-sample PVG. Moreover, the six-sample graph retained 82% of the all-sample PVG’s unique nodes and 81% of its edges (Table 3). Similarly, the three-sample PVG (151,760 bp; 150,557 bp without Ns) was 86 bp shorter than the six-sample PVG, and retained 94% of its nodes and 93% of its edges. The all- and six-sample PVGs contained 24% and 6% more SNPs than the three-sample PVG, respectively. Using linear mapping with Minimap2 as a baseline, mapping with Giraffe to one-, three-, six- and all-sample PVGs had 2.66, 3.48, 3.94 and 4.01 times the CPU runtime and 1.63, 0.92, 0.81 and 0.90 times the maximum RAM requirements, respectively. These results illustrated that the three- and six-sample PVGs captured most LSDV genomic diversity and required substantially less computational time for analyses than the all-sample PVG.

**Table 3.**
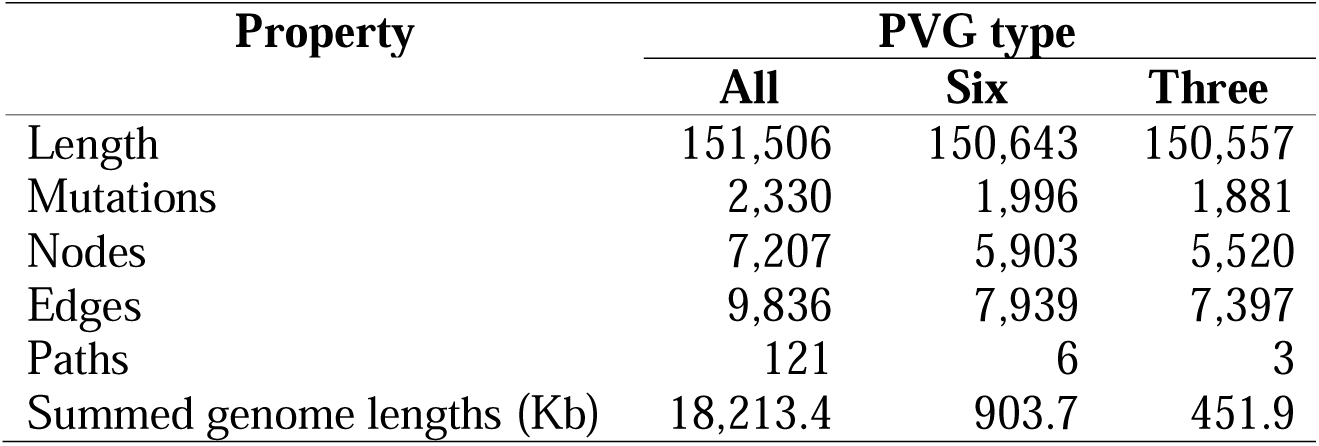
Properties of the all-sample (‘All’), six-sample (‘Six’), and three-sample (‘Three’) PVG. The length excludes bases denoted as “N”.

### Performance of PVG versus linear reference mapping across library types

We examined all available Illumina read libraries following careful quality control and viral read extraction (Table 4) to compare the outputs of mapping to these PVGs versus a single reference genome (Table S3, Supplementary Text 3). Reads were mapped to a linear reference (KX894508) with Minimap2, and to the one-, three-, six and all-sample PVGs using Giraffe. The one-sample PVG was composed of duplicate KX894508 sequences differing at the first base only. We examined the numbers of reads mapping in the resulting GAM and BAM files in the amplicon (n=27, Figure S4) and WGS libraries (n=46, Figure S5) and found that read libraries could be mapped well with either PVG-based or linear mapping approaches (Table S4). The differences between the median Minimap2 mapping rate and the four PVG options was negligible for amplicon libraries (all IQRs >99.5%). For WGS samples that had heterogeneous quality, Minimap2 had a slightly higher median mapping rate (73.6%) compared to the PVG options (medians 69.0%-73.0%). All six metagenomic libraries had mapping rates much lower overall with an IQR of just 17-35% based on linear read mapping and 16-32% for PVG mapping (Figure S6). Five of these libraries mapped better using linear reference mapping with Minimap2, with considerable heterogeneity across samples.

**Table 4.**
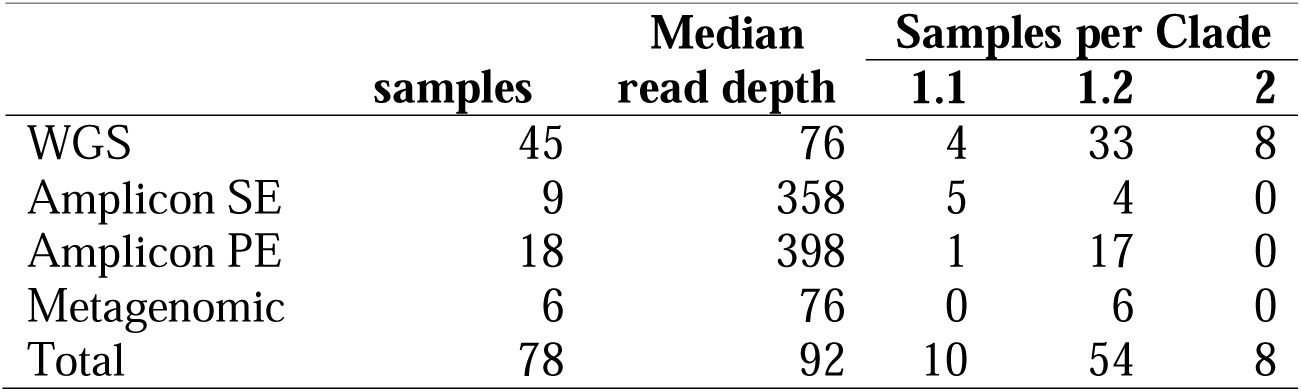
Properties of 78 valid LSDV read libraries examined. This excluded four mixed samples (SRR8442780, SRR19090746, SRR19090747, SRR19090748).

### Enhanced SNP detection using PVG-based Giraffe mapping

Giraffe produced equivalent or superior SNP detection to Minimap2 when mapping to the one-sample PVG (Supplementary Text 4, Table S5). SNPs occurring in multiple samples were less likely to be errors: these are non-singleton SNPs. Giraffe identified more of these shared SNPs that Minimap2 missed than vice versa (28 vs 18). Across the samples, there were 217 unique SNPs found by Giraffe only, versus 125 SNPs unique to Minimap2. Moreover, Giraffe did not miss any SNPs detected by Minimap2 and found more SNPs per sample (median 5, IQR 4-6) than Minimap2 (median 3, IQR 3-4) in the 21 amplicon libraries with <100 SNPs (Figure 3). Most SNPs in the amplicon samples with >1,000 SNPs were shared by both methods: unique SNPs constituted only 0.2-0.4% per sample.

**Figure 3.**
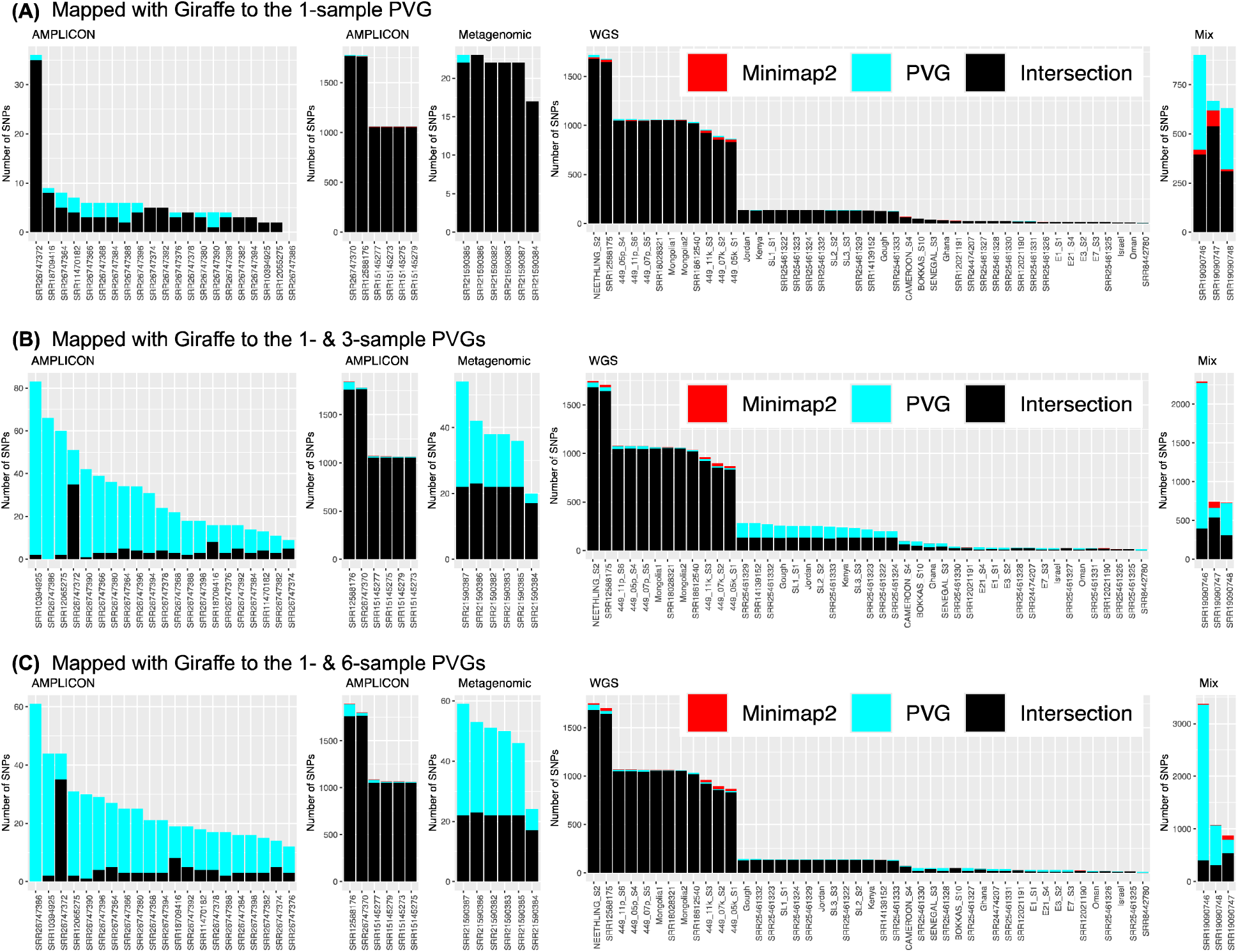
Number of SNPs detected by linear reference mapping with Minimap2 and Giraffe mapping to (A) one-sample PVG, (B) combined one- and three-sample PVGs and (C) combined one- and six-sample PVGs. Red (Minimap2) indicates SNPs captured by Minimap2 alone. Cyan (PVG) indicates those captured by PVG-based mapping only. Black (Intersection) indicates those captured by both Minimap2 and PVG mapping. Data is arrayed across library type: amplicon, metagenomic, WGS and mixed samples (Mix). The y-axes are different for each dataset, and those for amplicon datasets are split to match their corresponding SNP ranges. Three mixed samples (‘Mix’, right panel) are separate from the other WGS samples.

To accurately compare PVG-based versus linear mapping, we merged the detected SNPs from Giraffe mapping to both the one- and three-sample PVG, and the one- and six-sample PVG. This was essential to allow interpretation using a single genome coordinate system. The merged one- and three-sample SNP data captured far more non-singleton SNP sites missed by linear mapping than vice versa (353 vs 24 sites, Figure 3). This totalled 2,658 additional mutations across samples compared to 176 mutations uniquely associated with linear mapping (Table S7). Similar trends were also seen with the merged one- and six-sample SNP data dataset (Table S8). On average, ~267 SNPs/sample were shared between the two approaches, but more SNPs were unique to the merged PVG datasets (77-80 per sample) than to Minimap2 (~3 per sample) (Figure 4). In the WGS libraries, SNP detection was also higher in the merged PVG-based mapping (2,279 and 2,090 in the merged three- and six-sample PVG outputs, respectively) compared to Minimap2 (1,917). The same pattern was observed in amplicon, simulated, and metagenomic libraries. In addition, subconsensus SNP detection was higher for the PVG-based mapping approach (Figure S7, Figure S8, Supplementary Text 5). A median of 27% of SNPs from the three-sample PVG lacked corresponding sites for transfer to the linear reference KX894508, reflecting differences between reference sequence and observed states (Table S6). These SNPs were evenly spread among the genomes in the three-sample PVG (Figure 4A), but were unevenly shared in the genomes from the six-sample PVG, suggesting that the three additional genomes (OK318001, KX683219, MW732649) added limited extra information (Figure S9).

**Figure 4.**
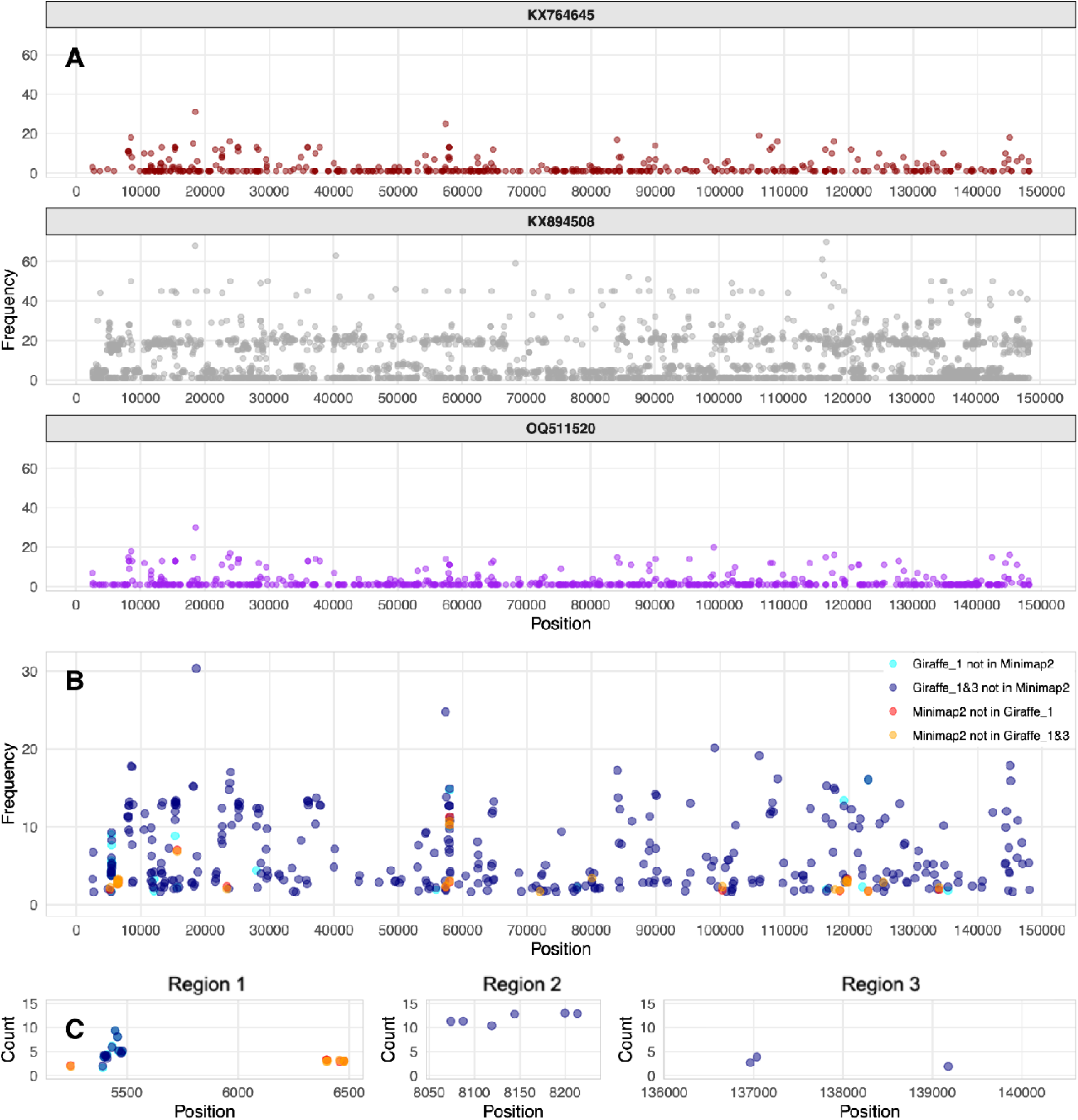
(A) Genome positions of mutations in the merged output of Giraffe mapping to the one- and three-sample PVGs showing the KX764645 (wine, top), KX894508 (grey, middle) and OQ511520 (purple, bottom) genomes. The bin size was 250 bp. (B/C) The number of novel non-singleton SNPs detected by different mapping approaches where (B) is genome-wide and (C) show Regions 1, 2 and 3. Cyan points are SNPs missed by linear mapping but found by Giraffe mapping to the one-sample PVG (28 SNPs genome-wide). Blue points are SNPs missed by linear mapping but found by Giraffe mapping to the merged one- and three-sample PVG output (352 SNPs genome-wide). Red points are SNPs missed by Giraffe mapping to the one-sample PVG (18 SNPs genome-wide). Orange points are SNPs missed by Giraffe mapping to the merged one- and three-sample PVG data (23 SNPs genome-wide).

We examined SNPs found in multiple samples at three regions with elevated variation: 5,200-6,500 bp (Region 1), 8,055-8,231 bp (Region 2) and 136,200-140,400 bp (Region 3) (Figure 4). Region 1 spanning LD008 (encoding a soluble interferon gamma receptor) and LD009 (encoding an alpha amanitin-sensitive protein) had seven SNPs at 5,831-5,897 bp (0.105 SNPs/Kb) (Figure S10). Region 2 spanning LD011 (encoding a CC chemokine receptor-like protein) had three SNPs at 8,062-8,131 bp (0.042 SNPs/Kb) (Figure S11). Region 3 spanning LD144 (encoding a kelch-like protein), LD145 (encoding an ankyrin repeat protein) and LD146 (encoding a phospholipase D-like protein) had five SNPs at 136,626-712 bp (0.057 SNPs/Kb) and three at 140,272-297 bp (0.11 SNPs/Kb) (Figure S12). In each case, Giraffe mapping to the combined one- and three-sample PVGs identified more non-singleton SNPs missed by Minimap2 than vice versa: 14 vs 2 at Region 1; 6 vs 0 at Region 2; and 3 vs 2 at Region 3 (Figure 5).

**Figure 5.**
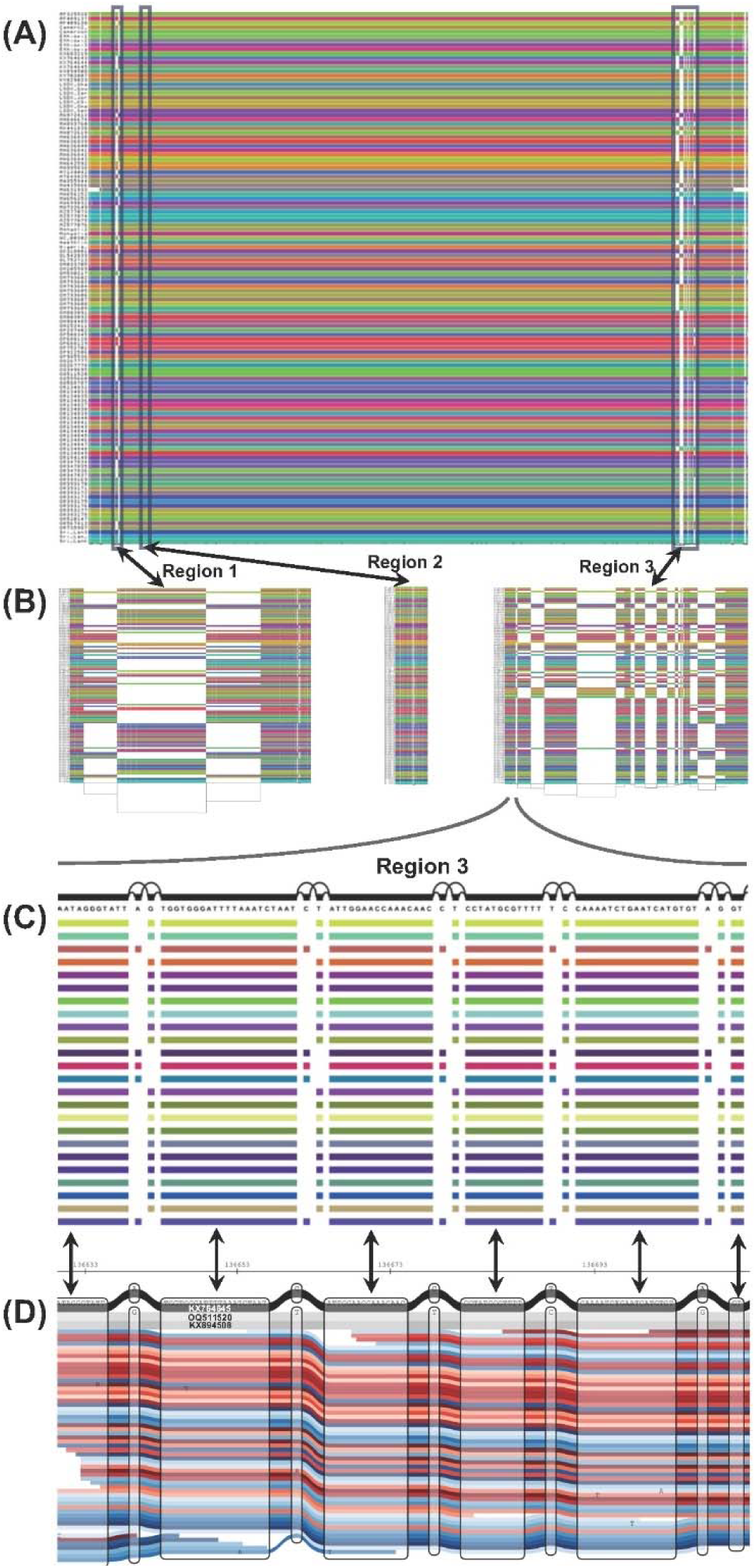
Visualisation of the LSDV PVG and the three identified regions of elevated variation where each panel shows increasing detail. (A) ODGI image of the complete PVG. (B) ODGI images of Region1-3, left: ‘Region1’ (5, 200-6,500 bp), middle; ‘Region2’ (8,055-8,231 bp), and right: ‘Region3’ (136,200-140,400 bp). (C) VG visualisation focussing on a region of LD144 (136,626-136,712 bp) within ‘Region3’, with five non-singleton SNPs indicated. The arrows between (C) and (D) denote the matching regions between these panels. (D) SequenceTubeMap image of reads from LSDV sample 449_07p_S5 in red and blue mapped to the three-sample PVG across the same region of LD144 (136,626-136,712 bp), showing the same five non-singleton SNPs, where KX894508 (Clade 1.2) is in light grey, OQ511520 (Clade 2) is in grey, and KX764645 (Clade 1.1) is coloured black. The nodes contain sequences, and the edges join these nodes across the reads.

We examined the numbers of SNPs detected in simulated read libraries for both OQ511520 (from Clade 2) and KX764645 (from Clade 1.1) compared to KX894508 as a function of read depth from 10-to 200-fold. This used the SNPs derived from genome alignment of OQ511520 versus KX894508 (1,130 SNPs) and KX764645 versus KX894508 (1,911 SNPs) as the truth set. Linear read mapping had better recall up to a read depth of ~90-fold, better precision, and resulted in a better predictive power based on the F1 score up to a depth of ~95-fold (Figure 6). Half of the real read libraries had a median depth < 95-fold. For read depths > 100-fold, the F1 score for PVG-based mapping was ~0.005 higher than linear mapping, which impacted functionally important variable regions. This was further illustrated by mapping the OQ511520 and KX764645 read libraries to each other’s references and comparing the SNPs within the OQ511520-KX764645 alignment as the true SNP set. PVG-based mapping identified all but one of 894 SNPs detected by linear mapping, and found 431 additional true SNPs that linear mapping missed (Figure S16). Furthermore, the additional SNPs detected using PVG-based methods alone in these simulated read libraries were at certain regions of high genetic diversity (Figure S13). For example, there were four nonsynonymous SNPs at the 3’ end of LD008 encoding a key immunomodulation protein: S236G (at 5,548 and 5,549 bp), N237S (at 5,551 bp), and D238V (at 5,554 bp) (Figure 4C). Another cluster of three nonsynonymous SNPs was found at the 3’ end of LD067 (encoding a host range protein, similar to VACV C7L): E183D (at 57,953 bp), E184D (at 57,965 bp) and D185E (at 57,959 bp) (Figure S14).

**Figure 6.**
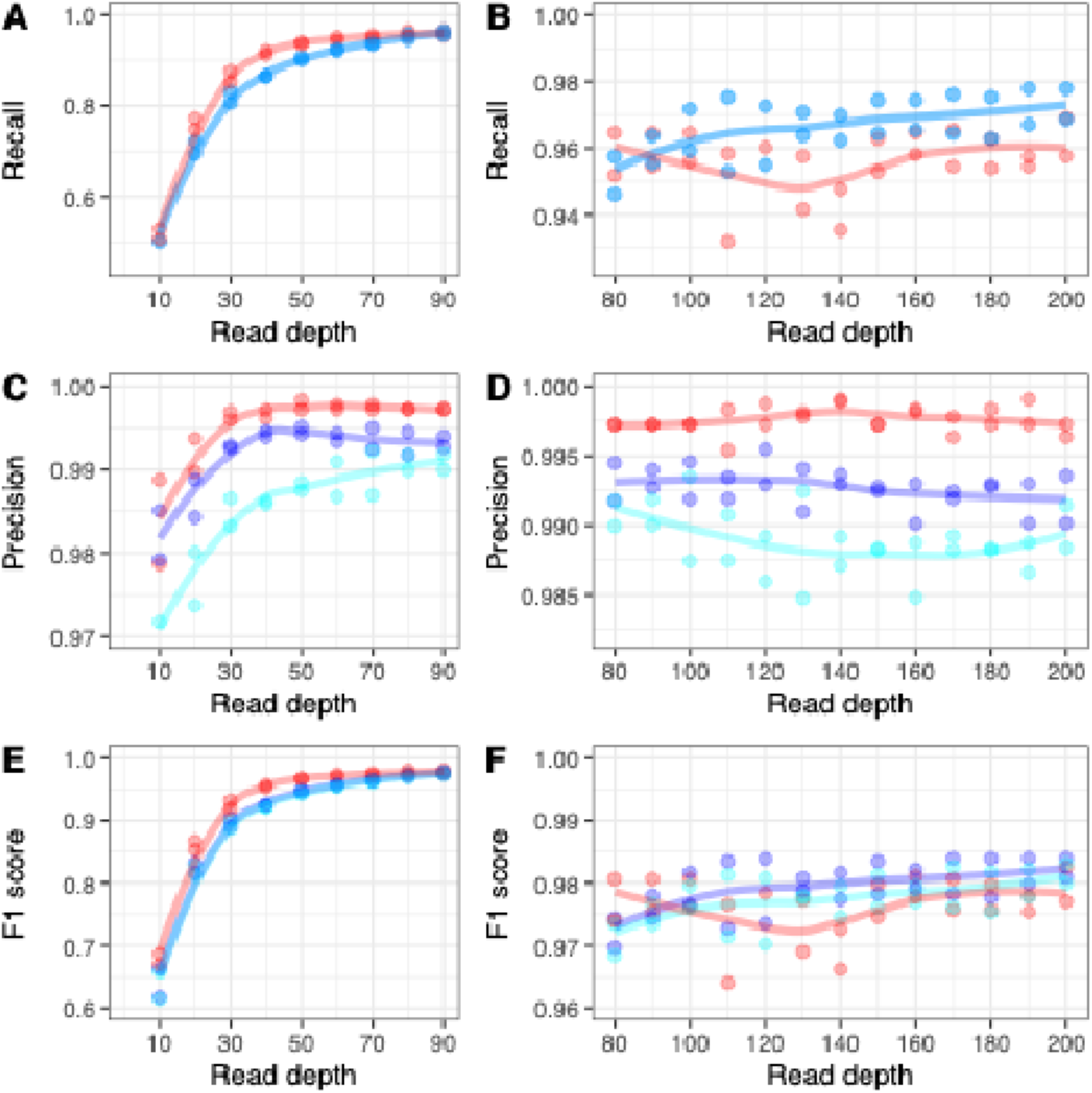
The effect of read depth on (A/B) recall, (C/D) precision, and (E/F) F1 score for genome-wide SNPs based on simulated reads for KX764645 and OQ511520. The red points and lines indicate patterns for linear read mapping with Minimap2. The blue points and lines indicate patterns for Giraffe mapping to the combined one- and three-sample PVG datasets. The cyan points and lines indicate patterns for Giraffe mapping to the combined one- and six-sample PVG datasets. The lines reflect the loess-smoothed linear trends in each dataset.

Furthermore, PVG-based mapping was more effective at SNP detection in two mixed samples: Lumpivax vaccine specimens previously reported as combinations of LSDV Clades 1.1, 1.2, GTPV and SPPV [86]. PVG-based mapping with Giraffe captured more within-sample diversity that linear read mapping with Minimap2 missed by mapping 13% and 26% more reads across these two samples (Supplementary Text 5, Figure S15).

### Phylogenetic and functional effects of novel SNPs from pangenome variation graph mapping

Phylogenies were constructed from 78 read libraries using SNPs from Minimap2 mapping to a linear reference or Giraffe to the combined one- and three-sample PVGs (Figure 7). The latter dataset resolved more population structure compared to mapping to a linear reference, particularly within Clade 1.2. Nonetheless, both identified several unique lineages with long external branches that were more basal, symptomatic of recombination and phylogenetic uncertainty. Using the same approach with SNPs ascertained with Giraffe mapping combined one- and six-sample PVGs showed no clear differentiation of groups within Clade 1.2, unlike Giraffe mapping combined one- and three-sample PVGs (Figure S17).

**Figure 7.**
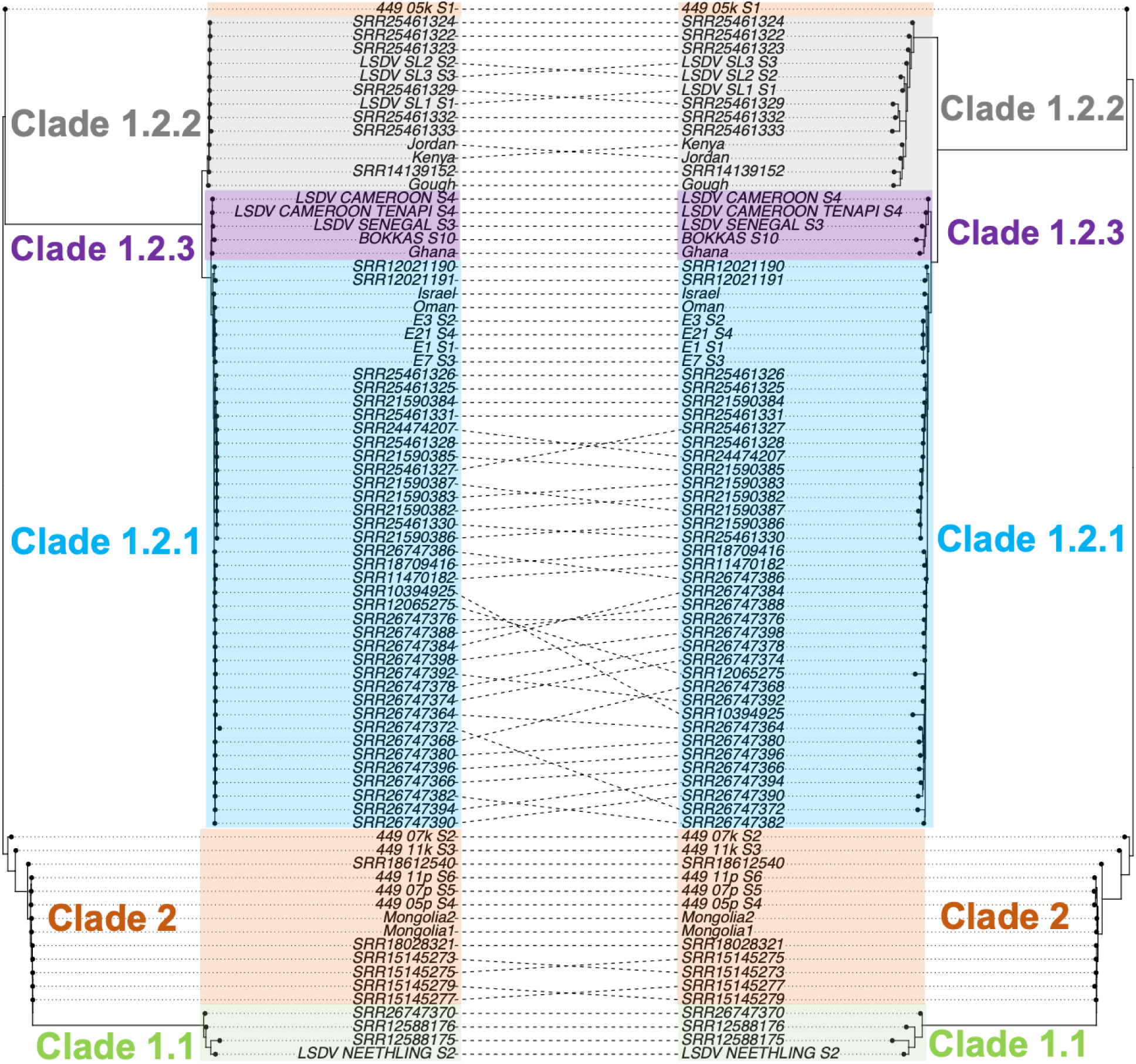
A co-phylogeny of genome-wide SNPs inferred from (left) linear reference mapping versus (right) PVG-based mapping. The clades are shown by shaded areas: Clade 1.1 in pale green, subclade 1.2.1 in pale cyan, subclade 1.2.2. in grey, subclade 1.2.3 in purple, and Clade 2 in pale orange.

Nonsynonymous (N) and synonymous (S) mutation rates from Minimap2 and Giraffe mapping to the one-sample PVG showed only subtle differences. Average N/S ratios were nearly identical (both 0.44), indicating strong consistency between approaches. Giraffe identified 0.3% more synonymous SNPs, 1.4% more nonsynonymous SNPs, and 2.4% more intergenic SNPs (Table 5). Both methods found that genes LD008, LD009, LD134, LD140 and LD145 had the highest SNP rates; these encode proteins associated with host recognition and immunomodulation. For example, the PVG-based approach uniquely identified E183D in LD067 (encoding a host-range protein) in all Clade 1.1 and 2 samples. A stop loss in LD019a (encoding a kelch-like protein) was detected in seven samples using PVG-based methods but linear mapping found it only three samples.

**Table 5.** Number of synonymous, nonsynonymous, intergenic, stop loss and stop gain SNPs detected in LSDV samples from Clades 1.1, 1.2 and 2. Linear reference mapping with Minimap2 (‘Minimap2’). Giraffe mapping to one-sample PVG (‘Giraffe_1’). Giraffe mapping to three-sample PVG (‘Giraffe_3’). Giraffe with merged one- and three-sample PVG (‘Giraffe_1&3’).

| Group | Minimap2 | Giraffe_1 | Giraffe_3 | Giraffe_1&3 |
| --- | --- | --- | --- | --- |
| **Synonymous** | 1,301 | 1,305 | 324 | 1,523 |
| **Nonsynonymous** | 581 | 589 | 418 | 894 |
| **Intergenic** | 84 | 86 | 46 | 117 |
| **Stop loss** | 3 | 3 | 4 | 5 |
| **Stop gain** | 1 | 2 | 32 | 34 |
| **Nonsyn/syn ratio** | 0.45 | 0.45 | 1.29 | 0.59 |

Giraffe mapping to the merged one- and three-sample PVGs recovered 17% more synonymous SNPs, 54% more nonsynonymous SNPs, and 39% more intergenic SNPs (Table 4) relative to Minimap2 mapping with a linear reference. This also highlighted additional potential functional changes. At LD012 (encoding an ankyrin repeat protein), G136E was in all subclades 1.2.2 and 1.2.3 samples, and an additional change (Q181*) was in subclade 1.2.2. At LD032 (encoding a polyA polymerase large subunit), S65G and K125* was in subclades 1.2.2 and 1.2.3. At LD135 (IFN-alpha/beta-binding protein), Y117*(128,683 bp) was in six isolates from Mongolia.

In total, 956 SNPs differentiated Clades 1.1 and 1.2 (Table 4). Ten genes accounted for 32% of these: LD134 (accounting for 66 SNPs), LD008 (52 SNPs), LD146 (39), LD009 (25), LD152 (23), LD039 (23), LD135 (22), LD049 (20), LD033 (19) and LD071 (18). Clade 2 showed less differentiation, consistent with its recent recombinant background, but carried three nonsynonymous changes absent in either Clade 1.1 or 1.2: N202D at LD008, N290D at LD059, and E96K at LD135. Within Clade 1.2, two nonsynonymous SNPs distinguished subclade 1.2.1 from 1.2.2/1.2.3, and four further SNPs separated 1.2.2 from 1.2.3.

## Discussion

In this study, we constructed and mapped reads to different LSDV PVGs to demonstrate that a small representative PVG captured nearly all known genomic diversity in the species. A PVG from only three assemblies, one per major lineage, recovered 97% of known SNP-level diversity while keeping graph size minimal. Comparisons with PVGs built from six and 121 genome assemblies (resulting in PVG extensions of 86 bp and 863 bp, respectively) showed only marginal gains in recovered variation, indicating that LSDV three-clade population structure provided an informative guide for selecting PVG representatives. The three-sample PVG retained 81% of all SNPs identified by the full 121-sample PVG. Mapping to PVGs had similar maximum RAM usage and at least 2.6 times the CPU time required using mapping to a linear reference as a baseline: previous work in both humans [1] and herpesviruses [10] indicated the compute require for both were more comparable.

Our comparisons between PVG-based and linear-reference mapping emphasised the limitations of relying on a single reference genome. Mapping to a PVG performed as well as or better than to mapping against a linear reference across diverse read libraries, consistent with previous analyses in other organisms [9]. Importantly, 27% of SNPs detected using PVGs could not be projected onto the linear reference, demonstrating the extent of reference bias and the inability of a single genome to represent the full spectrum of LSDV haplotypes [29]. PVG-based mapping also improved phylogenetic resolution: unlike linear mapping, it recovered variants differentiating subclades 1.2.1, 1.2.2 and 1.2.3. These gains are particularly relevant for detecting recombinant lineages containing combinations of wild-type and vaccine-derived ancestry, which continue to emerge across the LSDV geographic range [31–35].

Choosing tools to study viral diversity depends on whether the focus is within [versus between[sample variation. Within-sample relies on deep sequencing and highly sensitive callers to detect rare alleles, whereas between[host comparisons prioritise accurate representation of lineage[defining and structural variation across many genomes. Our simulated[data evaluation illustrated how these requirements influenced the relative performance of mapping strategies: PVG[based mapping yielded higher SNP accuracy if read depth was >95, consistent with its ability to reduce reference bias and resolve lineage[specific haplotypes. Linear[reference mapping was more reliable at depths <95. Furthermore, our work highlighted how PVG-based schema mapped far more reads from samples of mixed capripoxvirus origin. These results suggested that PVGs can complement existing tools by both supporting high[confidence detection of within[host mutations at sufficient depth and providing a unified, bias[reduced framework for broader population[wide analyses.

This implied a practical PVG design strategy for viruses with clear population structure to help moderate the challenge in interpreting complex datasets (Figure 8). We propose constructing two complementary PVGs: (i) a one-sample PVG representing the dominant clade, and (ii) a multi-sample PVG containing one high-quality genome per major lineage. Mutations identified from both PVGs can then be merged, enabling sensitive detection of mutations that would be overlooked using linear references alone. The number of genomes included in each PVG would be guided by lineage size, genetic uniqueness, and relevant epidemiological context.

**Figure 8.**
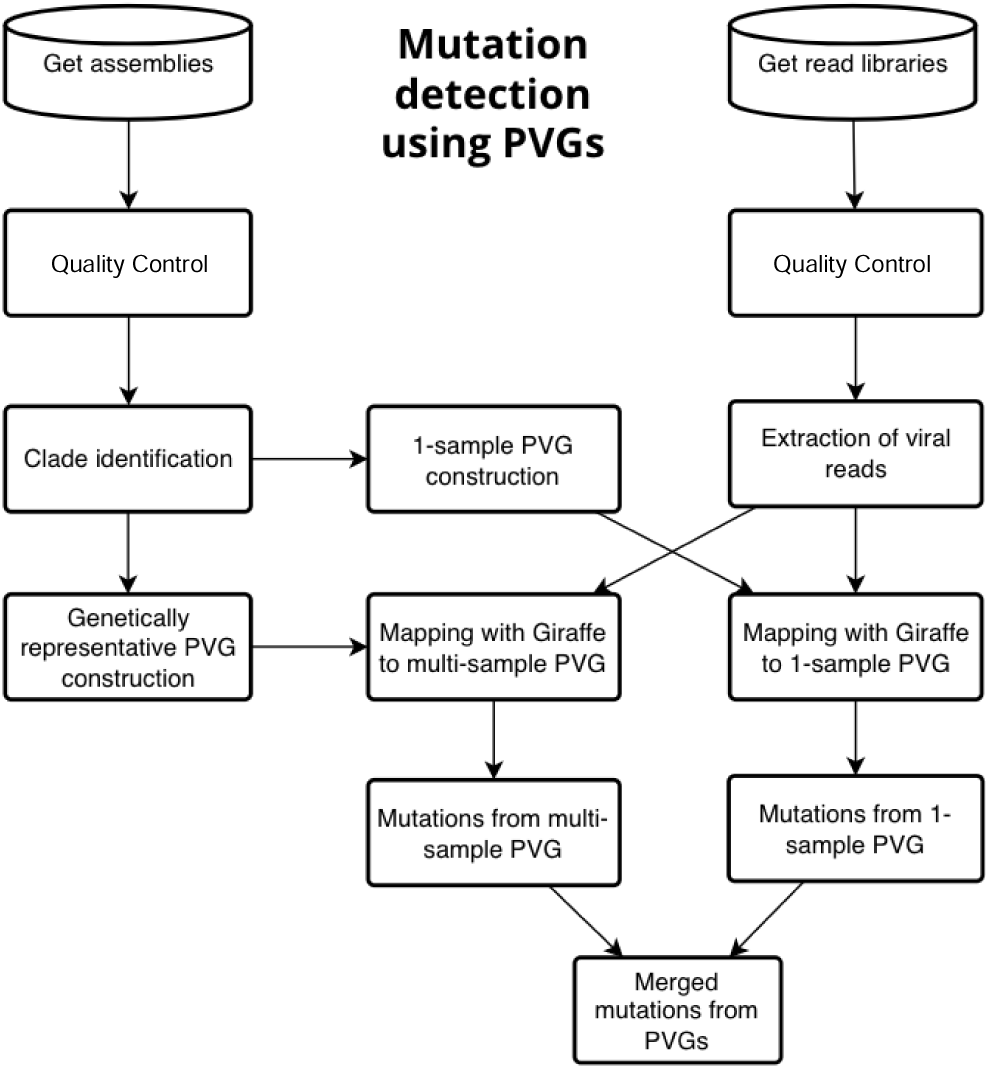
A flow chart for mutation detection using PVGs. Steps start from using genome assemblies to construct a one- and multi-sample PVG. Read libraries can then be mapped to both PVGs to discover mutations by merging the PVG-based outputs.

This study had five key limitations. First, newer datasets were generally higher quality, reflecting advances in technology. Secondly, the sequencing technology mattered: WGS read libraries were more heterogeneous, whereas amplicon ones were more consistent and benefited most from PVG-based mapping. The AT-rich LSDV genome (27% GC) [28] likely contributes these effects. Third, the diversity of samples across technologies was mixed: all eight Clade 2 read libraries were from WGS, and the six metagenomic samples were all from Clade 1.2. Fourth, although long-read assemblies are ideal for PVG construction because they better resolve structural variants and repetitive regions, some older Nanopore datasets had insufficient base quality for reliable inclusion. Fifth, our approach is well-suited to viruses with relatively simple population structures such as LSDV, highly diverse RNA viruses or within-host quasispecies may require more specialised PVG strategies.

Overall, we show that PVGs provide substantial advantages for analysing viral genomic diversity. For LSDV, a small lineage-representative PVG captured nearly all known variation, and PVG-based mapping with Giraffe identified more biologically meaningful mutations than linear reference mapping. LSDV and other poxviruses possess particular evolutionary features that made them well suited to PVG-based analyses: they have slow mutation rates, but higher rates of recombination and structural rearrangements generate substantial genomic diversity. Our framework offers a generalisable strategy for building viral PVGs and has direct implications for mixed-sample detection, genomic surveillance [87], outbreak reconstruction, and identifying recombinant vaccine-derived lineages in LSDV [88] and other DNA viruses.

## Abbreviations

ADE: automated de novo error
AF: allele frequency
BAM: Binary Alignment Map
BQ: base quality
CDS: coding sequence
GAM: Graph Alignment Map
GBWT: graph Burrows–Wheeler transform
GFA: Graphical Fragment Assembly
GFF: general feature format
GTF: gene transfer format
GTR: general time reversible
IGV: Integrative Genomics Viewer
IQR: interquartile range
Kb: kilobase
LD: Lumpy skin disease virus gene designation
LSDV: Lumpy skin disease virus
MAFFT: multiple alignment using fast Fourier transform
MQ: mapping quality
NCBI: National Center for Biotechnology Information
ODGI: Optimized Dynamic Genome/Graph Implementation
PAV: presence–absence variant
PCA: principal component analysis
PGGB: Pangenome Graph Builder
PVG: pangenome variation graph
QC: quality control
RDAF: read-depth allele frequency
SAM: Sequence Alignment Map
SNP: single nucleotide polymorphism
Ti/Tv: transition–transversion ratio
VCF: variant call format
VG: Variation Graph toolkit
WGS: whole-genome sequencing
wfmash: weighted fast minimizer-based mash-like aligner

## Supplementary Table Legends

Table S1. The Illumina read libraries’ sample names, assigned clade/subclade (Clade), library type (Type), read type (Layout), number of bases (bases), average length (avgLength), size in MB (size_MB), experiment ID (experiment), library name (LibraryName), library selection (LibrarySelection), sequencing machinte (Model), Bioproject ID, biosample ID, sample name and additional information (where relevant). Libraries 449_05k_S1, 449_05p_S4, 449_07k_S2, 449_07p_S5, 449_11k_S3 and 449_11p_S6 were newly sequenced. MeanReadDepth refers to the average read depth per sample across the genome from SAMtools.

Table S2. The quality control metrics for each read library, showing the numbers of reads pre-QC (before_filtering) and post-QC (after_filtering), the number of bases with BQ scores > 30 pre_QC (Q30_bases_before) and post-QC (Q30_bases_after), numbers of failing QC due to low quality (reads_failed_low_quality) or too many N bases (reads_failed_too_many_N) or if they were too short (reads_failed_too_short) or containing adapters (reads_with_adapter_trimmed) or partial adapters (bases_trimmed_due_to_adapters), and the duplicate read numbers (duplication_rate). Paired read libraries were merged so that each sample was represented by one set of read libraries.

Table S3. The number and percentage of reads associated with each read library across the major common name species found (cow, deer, sheep, human). Capripoxvirus reads are allocated under the columns “virus”.

Table S4. The reference name (rname), read depth, BQ and MQ across the read libraries organised by their mapping approach. The mapping approaches were Minimap2 to a linear reference, or Giraffe to the 1- or 3- or 6- or all-sample PVGs.

Table S5. Ti/Tv rates for different library types (amplicon and WGS) based on the mapping tool (Minimap2 vs Giraffe), PVG type (1-, 3-, 6-, all-sample or linear) and mutation caller (BCF stands for BCFtools, FB for Freebayes) for all SNPs and invalid ones.

Table S6. Fraction of valid SNPs that could not be lifted over from the sample’s coordinates to KX894508 if the allelic options allow. BCF stands for SNPs detected with BCFtools. FB stands for SNPs detected with Freebayes. VG stands for SNPs detected with vg. 3, 6 and ALL stand for mapping approaches using the 3-, 6- and all-sample PVGs, respectively. The mapping approaches using Giraffe are labelled GBWT.

Table S7. The numbers of SNPs discovered using Giraffe mapping to the combined one- and three-sample PVG Giraffe mapping compared to that for Minimap2 to a linear reference across samples. PVG denotes the number of SNPs found by PVG-based mapping. Minimap2 denotes the number of SNPs found by mapping to a linear reference. Total_all denotes the number of SNPs found by either option. Percent_PVG is the percentage of SNPs uniquely found by PVG-based methods, and Percent_M is the percentage of SNPs uniquely found by Minimap2. The samples are sorted by Library_type and then by the number of Minimap2 SNPs.

Table S8. The numbers of SNPs discovered using Giraffe mapping to the combined one- and six-sample PVG Giraffe mapping compared to that for Minimap2 to a linear reference across samples. PVG denotes the number of SNPs found by PVG-based mapping. Minimap2 denotes the number of SNPs found by mapping to a linear reference. Total_all denotes the number of SNPs found by either option. Percent_PVG is the percentage of SNPs uniquely found by PVG-based methods, and Percent_M is the percentage of SNPs uniquely found by Minimap2. The samples are sorted by Library_type and then by the number of Minimap2 SNPs.

## Acknowledgements

We acknowledge the Pirbright Institute’s Bioinformatics STP and Non-vesicular Reference Laboratory (NVRL).

## Author contributions

TD; Conceptualisation. CT, CW, LD, TD; Methodology. TD; Investigation. CT; Resources. TD; Data Curation. CW, LD, TD; Writing – Original Draft Preparation. CW, LD, TD; Writing – Review and Editing. CT, CW, LD, TD; Visualisation. TD; Project Administration.

