## Supplementary material for "Using pangenome variation graphs to improve mutation detection in a large DNA virus": Suppl_data

*Supplementary Data*

**Table of contents**

*Page Section*


1 1 Supplementary Text 1: Extensive host and contaminant DNA in LSDV read libraries
4 2 Supplementary Text 2: LSDV population structure and community identification
10 3 Supplementary Text 3: Mapping with Giraffe results in higher read depth & lower mapping quality
11 4 Supplementary Text 4: Varied mutation detection accuracy across library types, mapping tools & mutation callers
14 5 Supplementary Text 5: PVGs deliver better SNP detection in mixed samples
15 Figure S1
16 Figure S2
17 Figure S3
18 Figure S4
19 Figure S5
20 Figure S6
21 Figure S7
22 Figure S8
23 Figure S9
24 Figure S10
25 Figure S11
26 Figure S12
27 Figure S13
28 Figure S14
29 Figure S15
30 Figure S16
31 Figure S17

### 1 Supplementary Text 1: Extensive host and contaminant DNA in LSDV read libraries

Many samples had high rates of contamination in the form of reads with high similarity to the cow, deer, sheep, human and certain bacterial genomes (Figure S1.1). We examined the major mammalian contaminants (cow, deer, sheep, human) because these were prevalent across diverse samples, whereas bacterial or other contaminants were typically unique to specific samples. We performed host read removal by mapping each LSDV samples' reads against the following masked reference genomes with Bowtie2 v2.4.5 (Langmead & Salzberg 2012) (in the order provided) from Ensembl release 112 (April 2024): [1] *Bos taurus* (cow, Bos_taurus.ARS-UCD1.t3, GCA_002263795.3), [2] *Cervus hanglu yarkandensis* (Yarkand deer, Cervus_hanglu_yarkandensis.CEY_v1), [3] *Homo sapiens* (Homo_sapiens.GRCh38), and [4] *Ovis aries* (sheep, Ovis_aries.Oar_v3.1). We identified eight major bacterial species that were the most common non-mammalian contaminants identified by Kraken2: *Mesomycoplasma hyorhinis* strain IMT49388 (NZ_CP064323.1), *Metamycoplasma hominis* ATCC 27545 strain LBD-4 (NZ_CP009652.1), *Streptococcus iniae* strain DFSM220524 (NZ_CP125107.1), *Mannheimia varigena* USDA-ARS-USMARC-1296 (NZ_CP006943.1), *Moraxella lincolnii* strain 302605.3 (NZ_CP147511.1), *Arthrobacter* sp. TMP15 (NZ_CP154262.1), *Stenotrophomonas maltophilia* strain CGMCC 1.1788 (NZ_CP147720.1) and *Bacteroides xylanisolvens* strain CL11T00C41 (NZ_CP072212.1). The resulting contaminant and valid read libraries for each step were extracted using SAMtools v1.19.2 (Li et al 2009).

A number of observations were apparent in terms of minimising contamination (sensitivity): amplicon sequencing was much more effective, with median yields of >99.9% (Table S). WGS (median 53.5%) had a majority of reads that were LSDV, whereas metagenomic (median 1.6%) did not. The latter were mainly bovine (76.6%). It was also noticeable that the WGS yield per species (including LSDV) were highly variable relative to the other options. WGS libraries showed consistent low levels of human and deer reads, with higher levels of cow and sheep reads (Table S1.1).

Another goal of sequencing is to maximise the number of valid reads of interest (specificity) – here that was LSDV. Metagenomic (median 512k) and amplicon (medians 510k and 457k) sequencing were the most effective at this compared to WGS (median 219k). Again, WGS outcomes varied considerably. These results indicated that amplicon sequencing achieved the best balance of sensitivity and specificity, though with a considerable cost in the forms of design and staff time. Metagenomic approaches were excellent on specificity but poor on sensitivity, and WGS was intermediate for sensitivity and specificity.

We examined the effect of duplicate read removal in the six metagenomic libraries on SNP detection. These were all from Clade 1.2.1 and so were closely related to KX894508: this meant the total number of SNPs found by any method was low. We processed the deduplicated and non-deduplicated samples in the same way, just without SAMtools' markdup function for the non-deduplicated samples, focusing on BCFtools' output. Duplicate removal was important and supports a lower rate of false positive mutations (Table S1.2). Most additional SNPs were clear false positive changes related to polyA or polyT tracts.

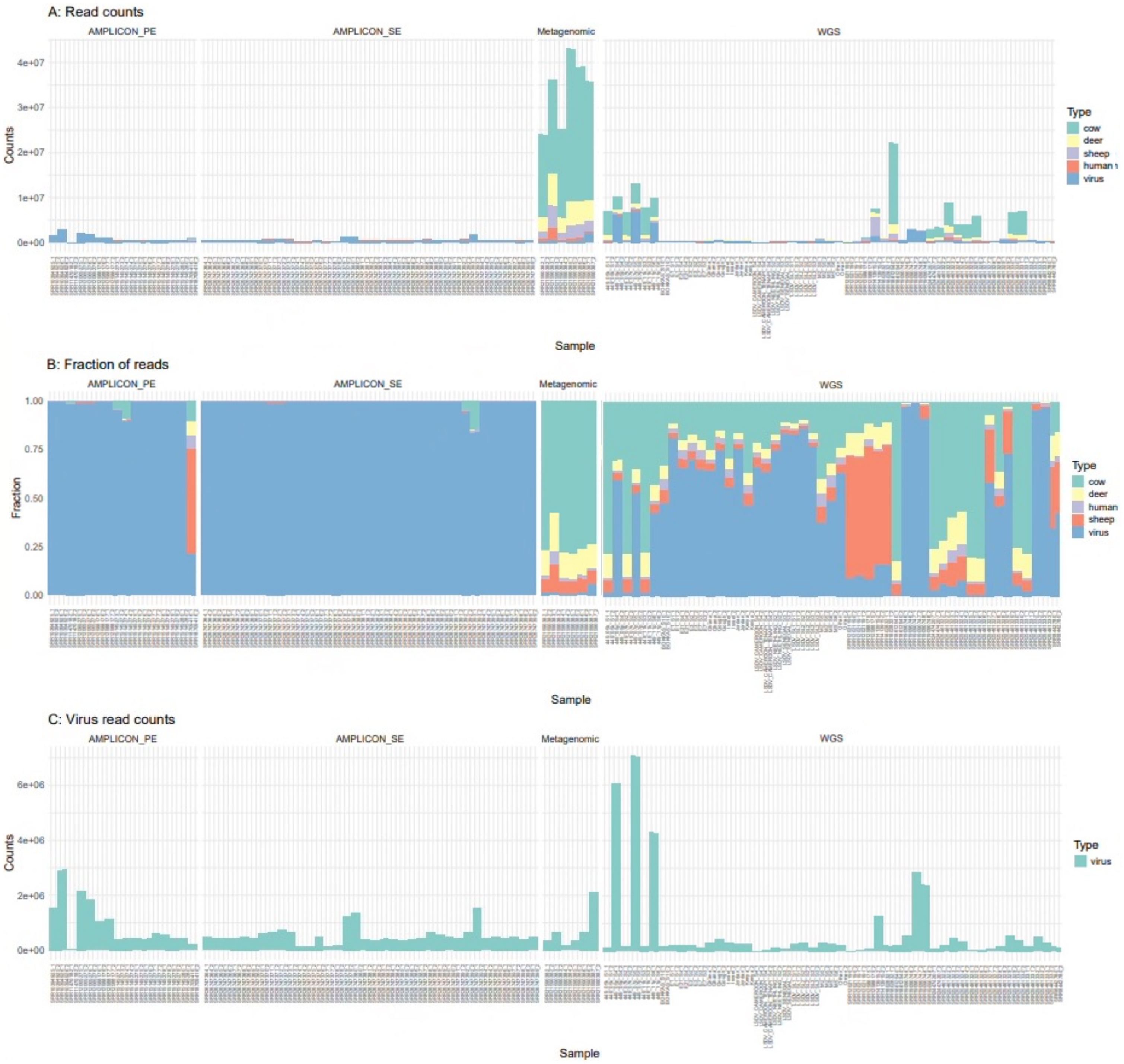


**Figure S1.1**. The reads (y-axis) per library matching different species (x-axis). The samples were organised by library type: amplicon PE, amplicon SE, metagenomic and WGS. (A) The numbers of reads per sample across all species: cow (green), deer (yellow), sheep (purple), human (red) and virus (LSDV, blue). (B) The fraction of reads per sample across all species. (C) The number of LSDV reads per sample (green). Note the y-axes scales differ between (A), (B) and (C).

| Library_type | Match | Median | SD | Median% |
|---|---|---|---|---|
| Amplicon PE | Cow | 284 | 26,261 | 0 |
| | Deer | 61 | 18,455 | 0 |
| | Sheep | 40 | 129,824 | 0 |
| | Human | 128 | 15,386 | 0 |
| | LSDV | 510,051 | 796,104 | 99.9 |
| Amplicon SE | Cow | 1 | 45,640 | 0 |
| | Deer | 0 | 510 | 0 |
| | Sheep | 0 | 495 | 0 |
| | Human | 8 | 2,646 | 0 |
| | LSDV | 457,180 | 299,367 | 100 |
| WGS | Cow | 104,769 | 3,118,132 | 21.3 |
| | Deer | 28,641 | 463,637 | 6.3 |
| | Sheep | 40,363 | 652,512 | 5.7 |
| | Human | 13,225 | 93,890 | 1.5 |
| | LSDV | 219,499 | 1,437,171 | 53.5 |
| Metagenomic | Cow | 23,700,629 | 5,891,283 | 76.6 |
| | Deer | 4,877,024 | 1,375,624 | 13.3 |
| | Sheep | 2,540,067 | 1,204,522 | 6.8 |
| | Human | 455,633 | 796,567 | 1.3 |
| | LSDV | 512,228 | 682,294 | 1.6 |

**Table S1.1**. The median (Median), standard deviation (SD) and median percentage (Median%) of reads per library type and species to which the reads were assigned.

| Method | Status | SRR21590382 | SRR21590383 | SRR21590384 | SRR21590385 | SRR21590386 | SRR21590387 |
|---|---|---|---|---|---|---|---|
| Linear | Deduplication | 22 | 22 | 17 | 22 | 23 | 22 |
| Linear | NonDeduplication | 41 | 40 | 37 | 41 | 44 | 38 |
| Giraffe_PVG1 | Deduplication | 22 | 22 | 17 | 22 | 23 | 22 |
| Giraffe_PVG1 | NonDeduplication | 42 | 44 | 38 | 42 | 45 | 41 |
| Giraffe_PVG13 | Deduplication | 33 | 33 | 17 | 31 | 37 | 38 |
| Giraffe_PVG13 | NonDeduplication | 54 | 56 | 49 | 53 | 61 | 54 |
| Giraffe_PVG16 | Deduplication | 46 | 45 | 17 | 39 | 48 | 54 |
| Giraffe_PVG16 | NonDeduplication | 78 | 89 | 69 | 76 | 99 | 75 |

**Table S1.2.** The numbers of valid SNPs detected in deduplicated and non-deduplicated metagenomic read libraries.

### 2 Supplementary Text 2: LSDV population structure and community identification

Our dataset indicated three main Clades. Clade 1.2's geographic spread was across Europe, the Indian subcontinent, the Middle East and east Africa (Figure S2). Clade 1.1's samples were from southern Africa, except for MG872412 (Croatia, 2016) and OR134848 (Serbia, 2016). Clade 2 originated radiated recently in central and east Asia. There were five samples from four rarer recombinant lineages (labelled R here). Both R_2.4 samples (OM530217 and OR194148, both Russia, 2019) were the most similar to Clade 1.1, followed by R_2.1 (OL542833) and then R_2.6 (OR194148, Russia, 2019). R_2.2 (MT134042) (both Russia, 2019) was more closely related to Clade 1.2. This high rate of recombinant lineages associated with isolation in Russia is well-established (Sprygin et al 2018). There was some variation in the main clades' genetic distances at the core (Figure S2.1), 5' end (Figure S2.2) and 3' end (Figure S2.3) regions such that Clade 2 was more related to Clade 1.1 at the 5' region compared to the core and 5' end regions. This was supported by PCA visualisations (Figure S2.4).

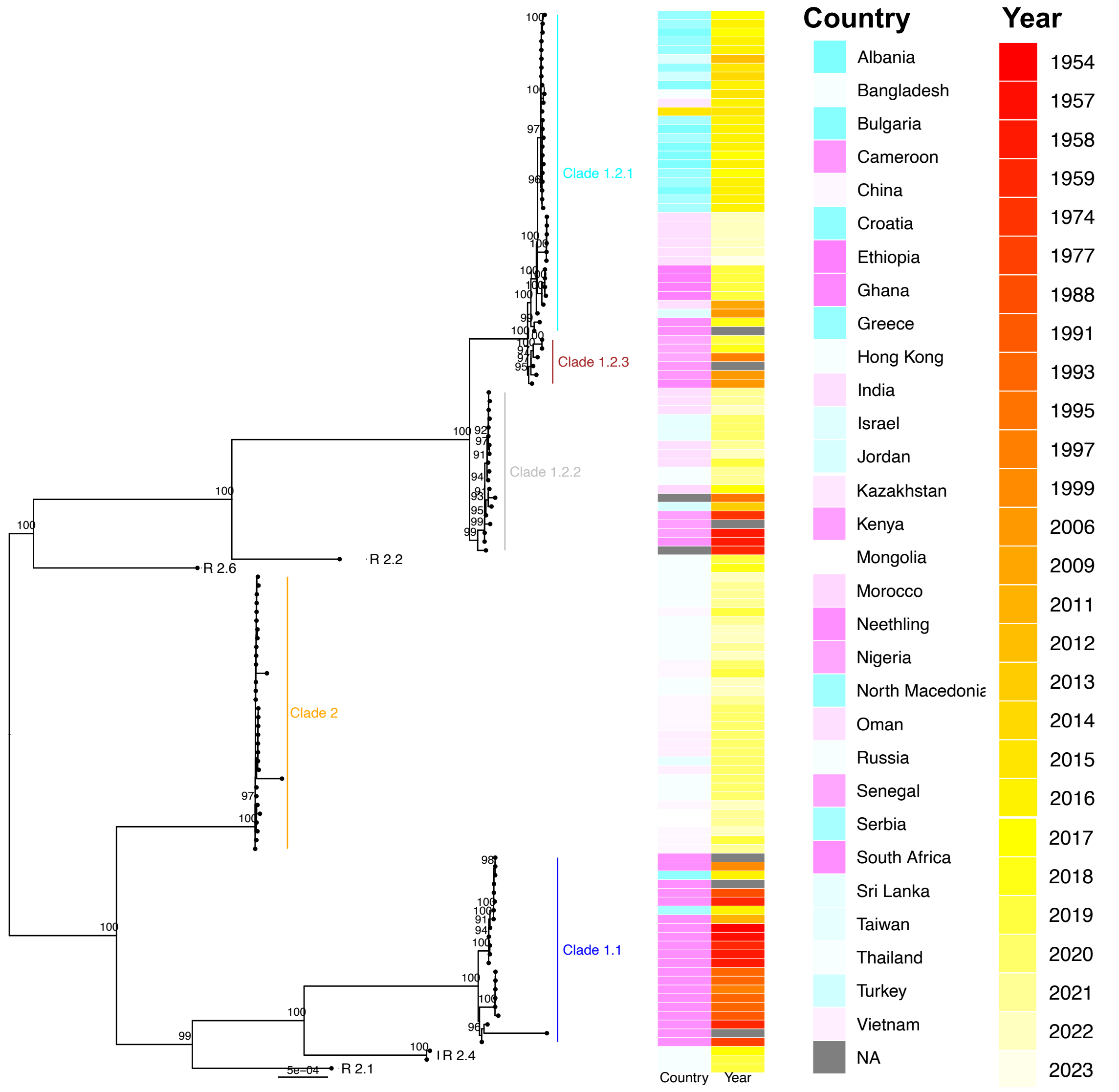


**Figure S2.1.** A phylogeny constructed with RAxML using a GTR+G4 substitution model based on SNP data for the core region (93 Kb). Clade 1.1 is highlighted in navy, Clade 1.2.1 in cyan, Clade 1.2.2 in grey, Clade 1.2.3 in brown and Clade 2 in orange. Rare recombinant (R) lineages are denoted 2.1, 2.2, 2.4 and 2.6. Right: the countries are show in the first column, and the year of isolation in the second column. Nodes with bootstrap support > 90 are shown. As expected, both R_2.4 samples (OM530217 and OR194148, both Russia, 2019) were most similar to Clade 1.1, followed by R_2.1 (OL542833) and R_2.2 (MT134042) (both Russia, 2019) was closely related to Clade 1.2. R_2.6 (OR194148, Russia, 2019) was equidistant from Clade 1.1 and 1.2.

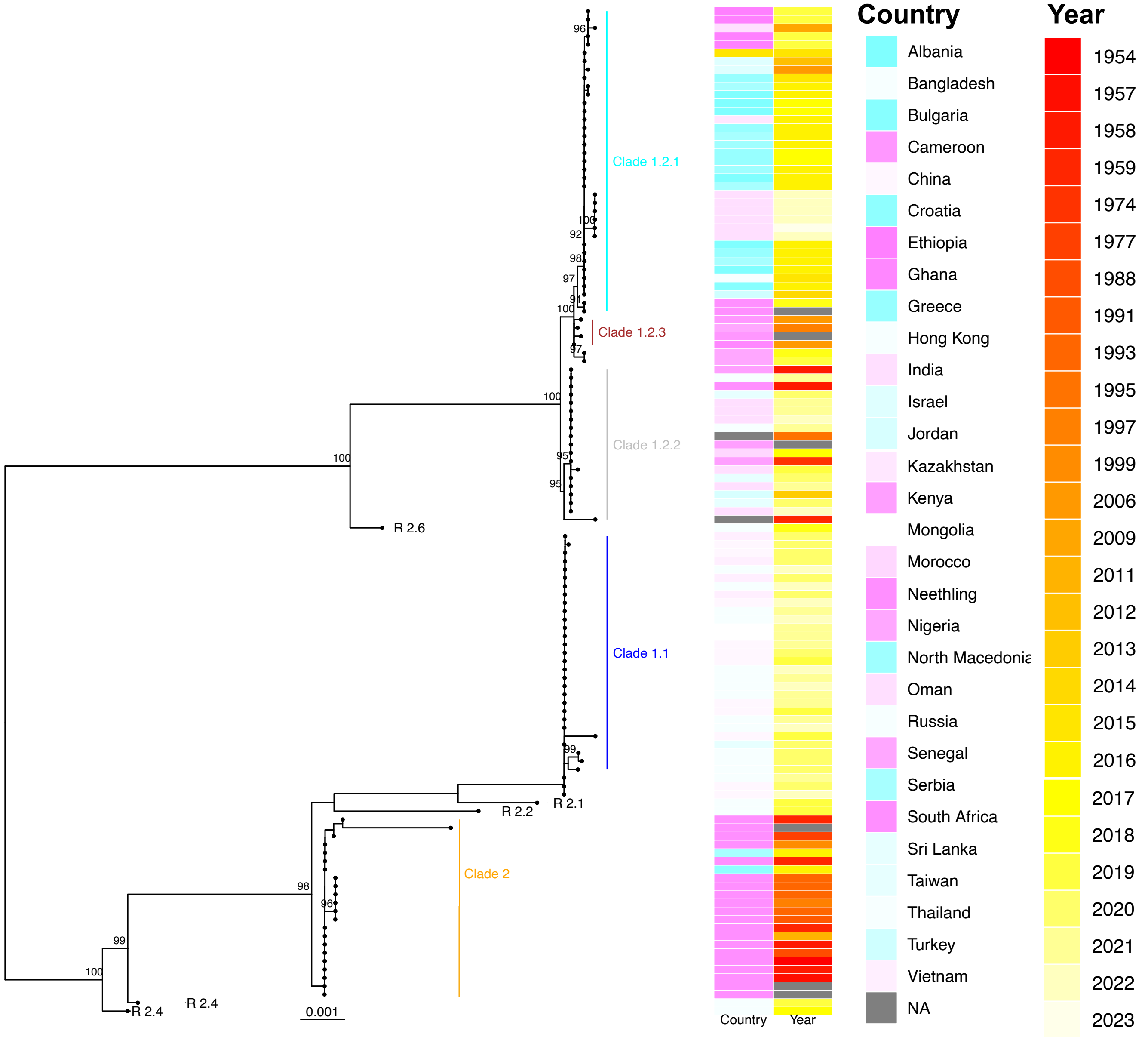


**Figure S2.2**. A phylogeny constructed with RAxML using a GTR+G4 substitution model based on SNP data for the 5' genome end. Clade 1.1 is highlighted in navy, Clade 1.2.1 in cyan, Clade 1.2.2 in grey, Clade 1.2.3 in brown and Clade 2 in orange. Rare recombinant (R) lineages are denoted 2.1, 2.2, 2.4 and 2.6. Clade 2 was more related to Clade 1.1 at the 5' genome end. Right: the countries are show in the first column, and the year of isolation in the second column. Nodes with bootstrap support > 90 are shown. R_2.1 (OL542833) was closely related to Clade 1.1. In contrast with the genome-wide SNPs, R_2.2 (MT134042) (both Russia, 2019) was closer to Clade 1.1; both R_2.4 samples (OM530217 and OR194148, both Russia, 2019) were not closely related to either Clades 1.1 or 1.2; and additionally, R_2.6 (OR194148, Russia, 2019) clustered closer to Clade 1.2.

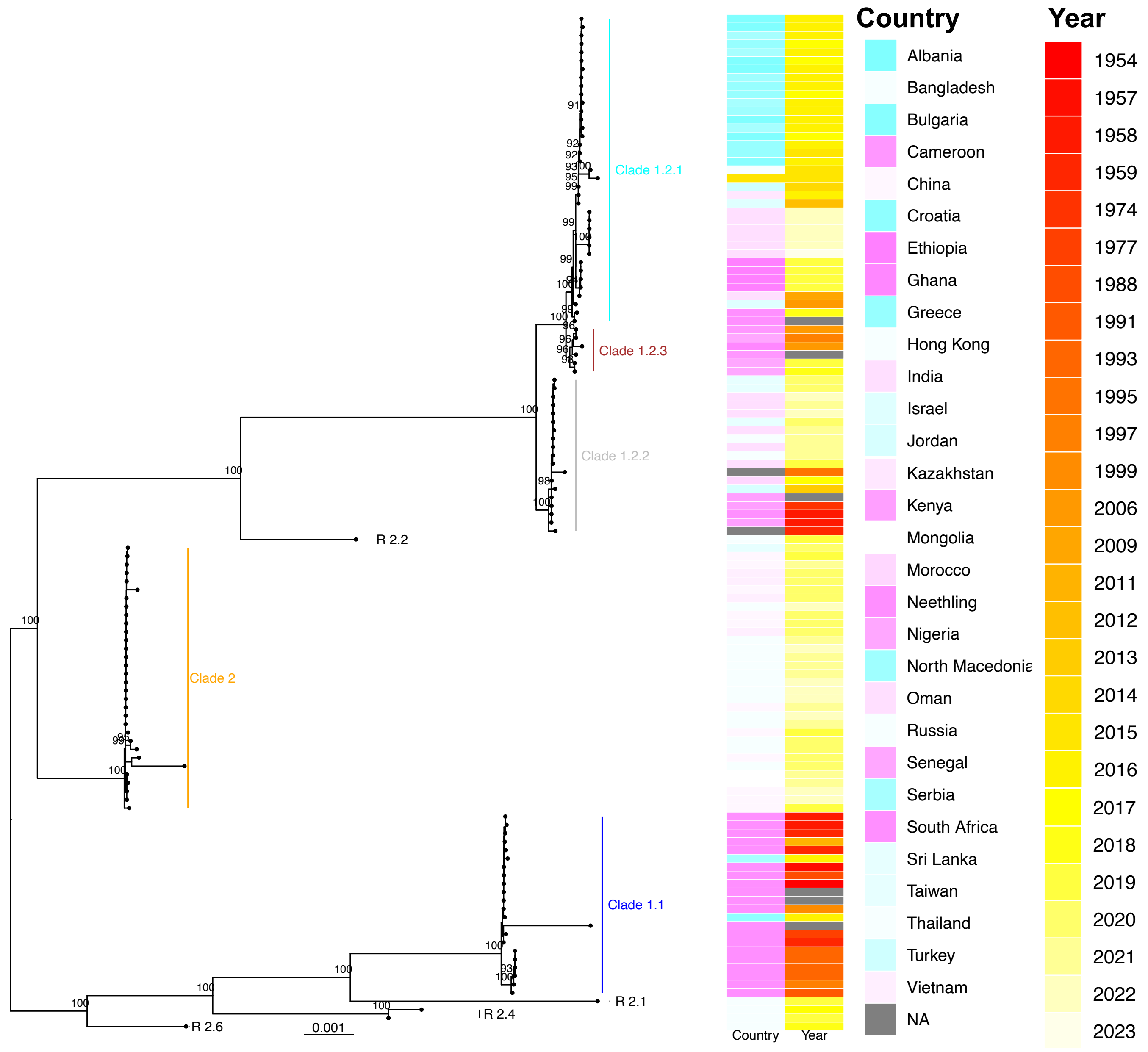


**Figure S2.3.** A phylogeny constructed with RAxML using a GTR+G4 substitution model based on SNP data for the 3' genome. Clade 1.1 is highlighted in navy, Clade 1.2.1 in cyan, Clade 1.2.2 in grey, Clade 1.2.3 in brown and Clade 2.2 in orange. Rare recombinant (R) lineages are denoted 2.1, 2.2, 2.4 and 2.6. Right: the countries are show in the first column, and the year of isolation in the second column. Nodes with bootstrap support > 90 are shown. In line with the genome-wide picture, R_2.1 (OL542833) was most similar to Clade 1.1, followed by both R_2.4 samples (OM530217 and OR194148, both Russia, 2019) and then R_2.6 (OR194148, Russia, 2019). As expected, R_2.2 (MT134042) (both Russia, 2019) was closely related to Clade 1.2.

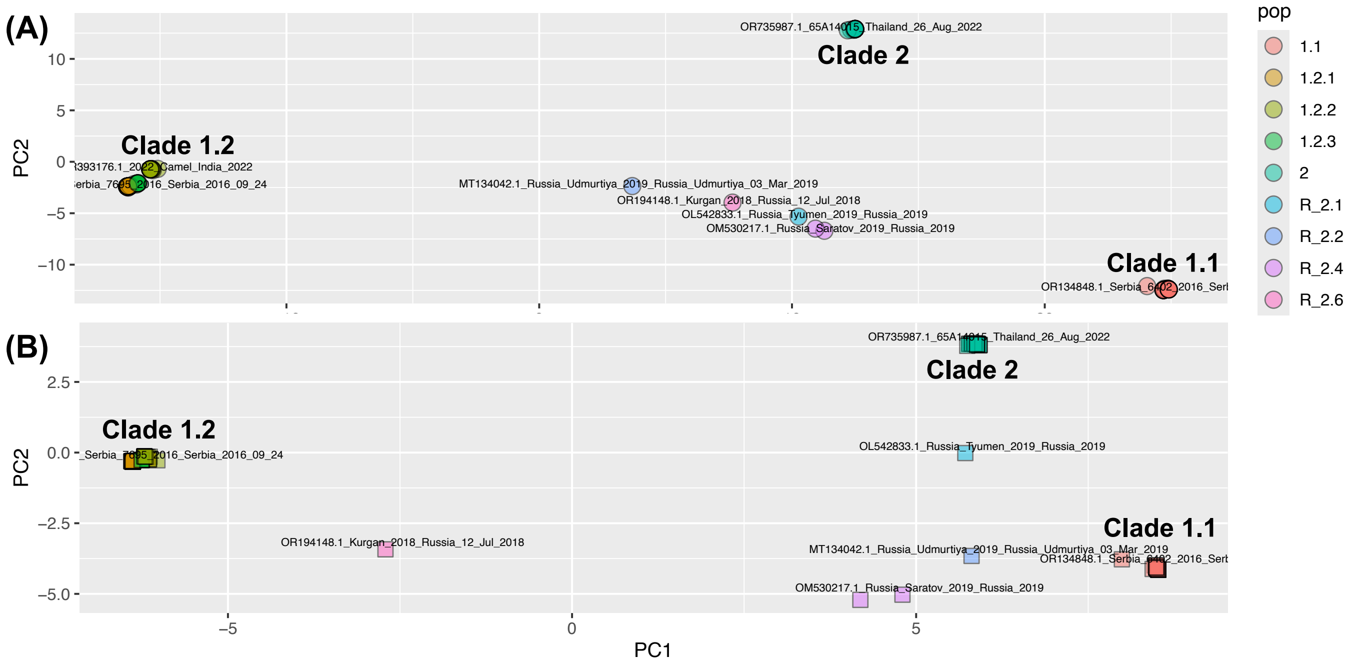


**Figure S2.4**. Principal components analysis (PCA) of the LSDV SNPs (A) genome-wide and (B) at the 5' end. (A) For genome-wide SNPs, PC1 had 71% of variation and PC2 had 19%. The LSDV phylogenetically defined Clades ("pop") are represented as 1.1 (red), 1.2.1 (light brown), 1.2.2 (brown-green), 1.2.3 (green) and 2 (cyan), and the rarer recombinants are shown as MT134042 (R_2.2, navy), OR194148 (R_2.6, mauve), OL542833 (R_2.1, blue), OM530217 (R_2.4, purple), and OR194148 (R_2.4, purple). (B) For the 5' end SNPs, PC1 had 79% of variation and PC2 had 14%. Here, both R_2.4 samples (OM530217 and OR194148, both Russia, 2019) were most like Clade 1.1, followed by R_2.2 (MT134042) (both Russia, 2019) which was genetically distant to Clade 1.1 at the core genome and 3' end. R_2.1 (OL542833) was more closely related to Clade 2. R_2.6 (OR194148, Russia, 2019) was more related to clade 1.2. Clade 2.

In addition, we found five major genetic groups (aka communities) in the 121 LSDV samples based on the PVG variation, corresponding to Clades 1.1, 1.2.1, 1.2.2, 2 and a subgroup of 1.1 (Figure S2.5). In the resulting network, R_2.1 (OL542833, Russia, 2019) was central, and was connected to Clades 1.1, 1.2.1, 1.2.2 and 2, reflecting its recombinant origins. It was connected to OM796304 and MW435866 from Clade 1.1. R_2.4 (OM530217, Russia, 2019 and MH646674, Russia, 2017), R_2.2 (MT134042, Russia, 2019) and R_2.6 (OR194148, Russia, 2019) clustered at the boundary between Clades 1.1 and 2, along with OP985536 (China, 2020) from Clade 2. The Clade 1.1 subgroup was composed of six samples from southern Africa and was also a subgroup in the whole-genome phylogeny (Figure S2). Patterns at the 5' (Figure S2.6), core (Figure S2.7) and 3' (Figure S2.7) and regions varied somewhat.

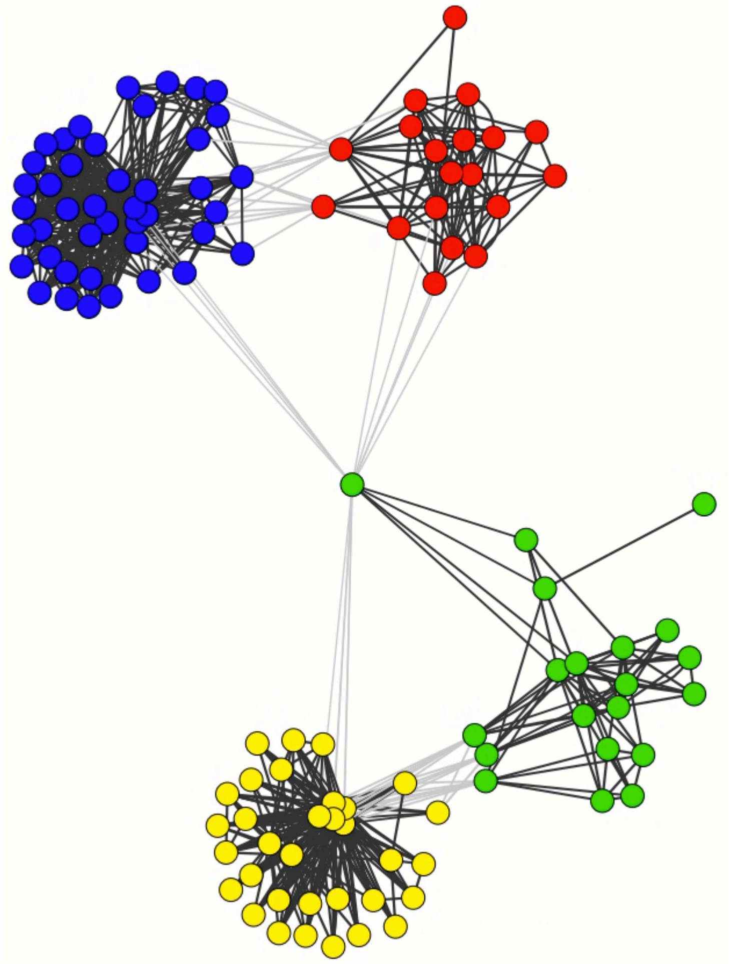

**Figure S2.5.** A network from genome-wide PVG data identified five groups corresponding to Clades 1.1 (green), 1.2.2 (red), 1.2.1 (blue), 1.1 (yellow) and a 1.1 subgroup (mauve) – the latter was genetically distinct, unlike the others. Clade 1.2.3 was within Clade 1.2.1 and is not coloured differently. Each node represents one sample. Grey lines indicate inter-community connections, and black lines indicate intra-community ones. OL542833 (R_2.1, Russia, 2019) had a central position in Clade 1.1 connected to Clades 1.2.1, 1.2.2 and 2. Both R_2.4 samples were allocated to separate clades here. OM530217 (R_2.4, Russia, 2019) and OR194148, (R_2.6, Russia, 2019) clustered with Clade 1.1 and were related to MH646674 (Russia, 2017) and other nodes in Clade 2. MT134042 (Russia, R_2.2, 2019) was related to diverse nodes in Clade 2, including OP985536 (China, 2020). In Clade 1.1 (green), OM796304 was related to OL542833, Neethling sample MW435866, and a divergent Pirbright Neethling sample. In Clade 1.2.2 (red), Gough (1959) was linked to samples from MU937760 (Russia, 2015) & Clade 1.2.3 (west Africa) and OP297402 (India, 2019) were related to samples from Clade 1.2.3.

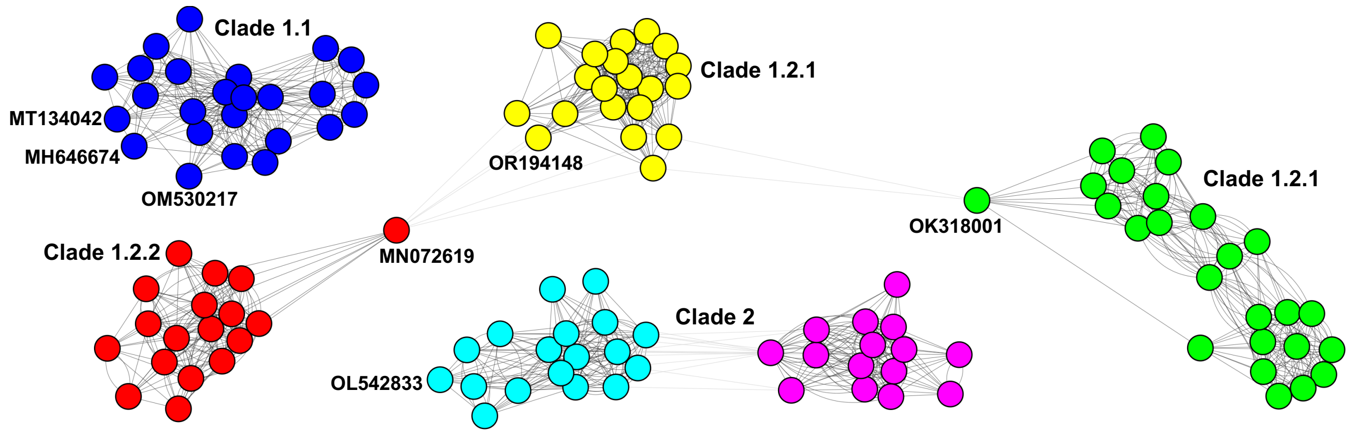


**Figure S2.6**. A network for the 5' PVG region (1-14 Kb) identified six communities corresponding to Clade 1.2.1 (green and yellow), Clade 1.2.2 (red), Clade 1.1 (blue), and Clade 2 (cyan and mauve). Clade 1.2.3 was within Clade 1.2.1 and is not coloured differently here. Clades 1.1 and 2 were genetically distinct, as was Clade 1.2. Each node represents one sample. Grey lines indicate inter-community connections, and black lines indicate intra-community connections. MN072619 (Kenya) was in 1.2.2 (red) but was connected to 1.2.1 (yellow). OK318001 (V281, Nigeria 2018) was in the green 1.2.1 cluster and was connected to the other yellow 1.2.1 group. OL542833 (R_2.1, Russia, 2019) was allocated to the cyan Clade 2 group. OM530217 (R_2.4, Russia, 2019), MH646674 (R_2.4, Russia, 2017) and MT134042 (Russia, R_2.2, 2019) were in the Clade 1.1 (blue) group. OR194148, (R_2.6, Russia, 2019) was in the yellow 1.2.1 group.

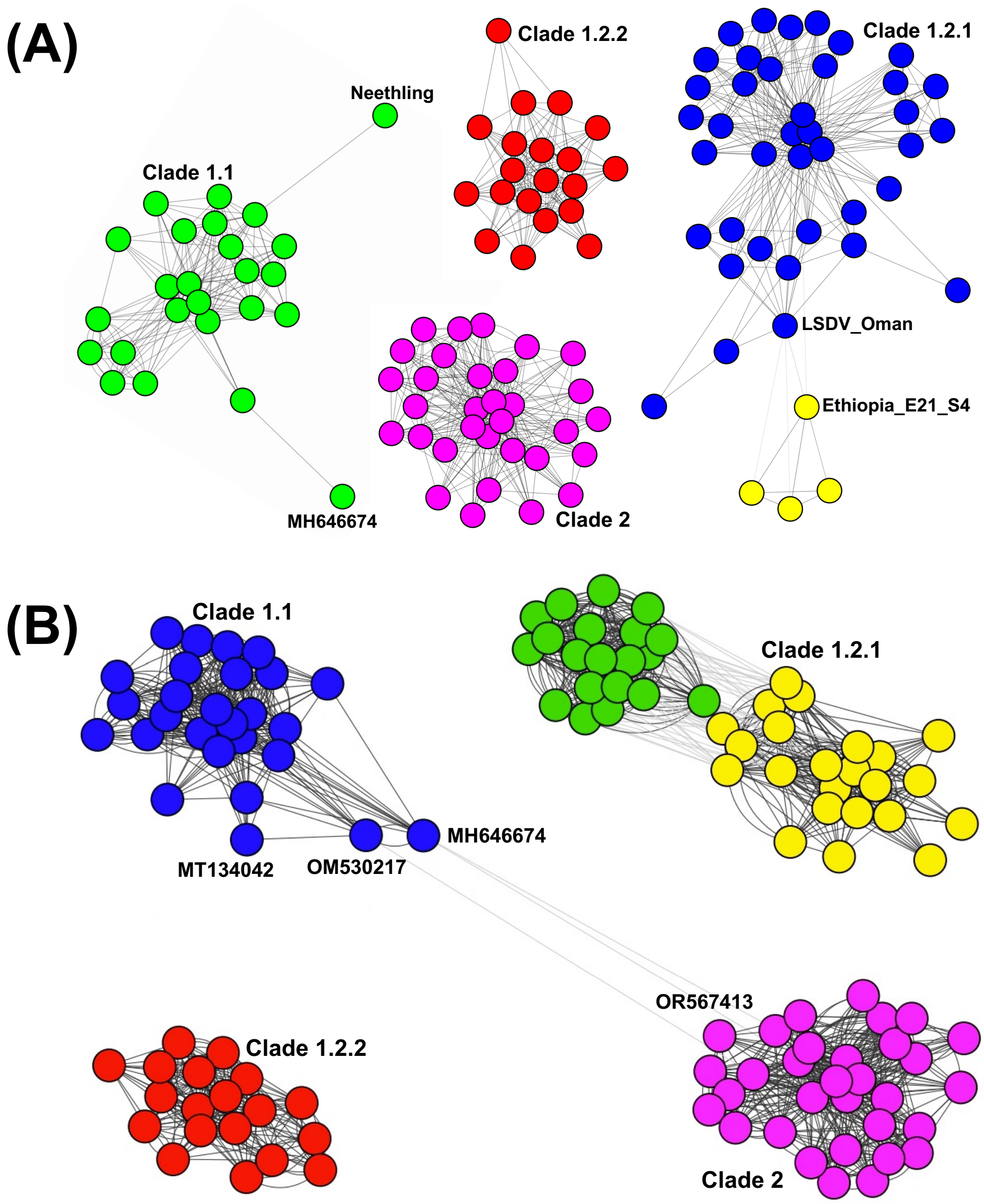


**Figure S2.7**. A network for the (A) core PVG region (14-106 Kb) and (B) 3' PVG end (106-148 Kb). Each node represents one sample. Grey lines indicate inter-community connections, and black lines indicate intra-community connections. Both identified five main communities including in (A), Clade 1.2.1 (green and yellow), Clade 1.2.2 (red), Clade 1.1 (blue), and Clade 2 (mauve). Clade 1.2.3 was within Clade 1.2.1 and is not coloured differently here; and in (B), Clade 1.2.1 (blue) connected to a subclade (yellow) comprised solely of samples from Ethiopia, where the other clades are 1.2.2 (red), 1.1 (green) and 2 (mauve). In (A), Clades 1.2.2 and 1.2.2 were genetically distinct, whereas Clades 1.1 and 2 were connected. In (A), MH646674 (R_2.4, Russia, 2017) and OM530217 (R_2.4, Russia, 2019) connected Clade 1.1 (blue) to OR567413 (China, 2022) and other samples in Clade 2 (mauve). MT134042 (Russia, R_2.2, 2019), OR194148 (R_2.6, Russia, 2019) and OL542833 (R_2.1, Russia, 2019) were allocated to the Clade 1.1 (blue). In (B), Clades 1.1, 1.2.1, 1.2.2 and 2 were genetically distinct. The Ethiopian subclade (yellow) was connected from Ethiopia_E21_S4 to 1.2.1 (blue) mainly by the sample LSDV_Oman (2009). The two outliers in Clade 1.1 (green) were MH646674.1 (R_2.4, Russia, 2017) and the TPI Neethling isolate. OM530217 (R_2.4, Russia, 2019) and OR194148, (R_2.6, Russia, 2019) were in the same group, 1.1. MT134042 (Russia, R_2.2, 2019) was in Clade 1.2.2 (red). OL542833 (R_2.1, Russia, 2019) is not shown in (B) because it had no high-confidence mappings with the other samples.

### 3 Supplementary Text 3: Mapping with Giraffe results in higher read depth & lower mapping quality

Although Giraffe mapped more reads than Minimap2 and VG-MAP, this resulted in a lower median MQ (Table S3.1). This suggested Giraffe mapping to the one-, three- and six-sample PVGs were viable options, whereas mapping to the all-sample PVG was not. When using the one-sample PVG, Giraffe mapped a median of 3.2-fold more reads than Minimap2, which increased to 4.5-fold when the three- or six-sample PVG was used. The median MQ values for Giraffe mapping to the one-, three- and six-sample PVGs were comparable to Minimap2's rate, but fell markedly for the all-sample PVG (Figure S3.1). This suggested the relative mapping probability increased as the allelic options in the PVG increased, but only up to a certain degree as evidenced by the poor all-sample PVG metrics. VG-MAP also had lower BQ and MQ negatively correlated with the number of samples in the PVG.

| Mapper | Reference | Depth | MQ |
|---|---|---|---|
| Minimap2 | Linear | 137 | 58.7 |
| Giraffe | one-sample PVG | 438 | 52.1 |
| Giraffe | three-sample PVG | 618 | 56.0 |
| Giraffe | six-sample PVG | 618 | 54.7 |
| Giraffe | all-sample PVG | 621 | 0.5 |
| VG-MAP | one-sample PVG | 138 | 58.7 |
| VG-MAP | three-sample PVG | 141 | 17.5 |
| VG-MAP | six-sample PVG | 141 | 16.9 |
| VG-MAP | all-sample PVG | 114 | 3.3 |

**Table S3.1**. The median depth and mapping quality (MQ) values for each mapper (Minimap2, Giraffe, VG-MAP) and reference set.

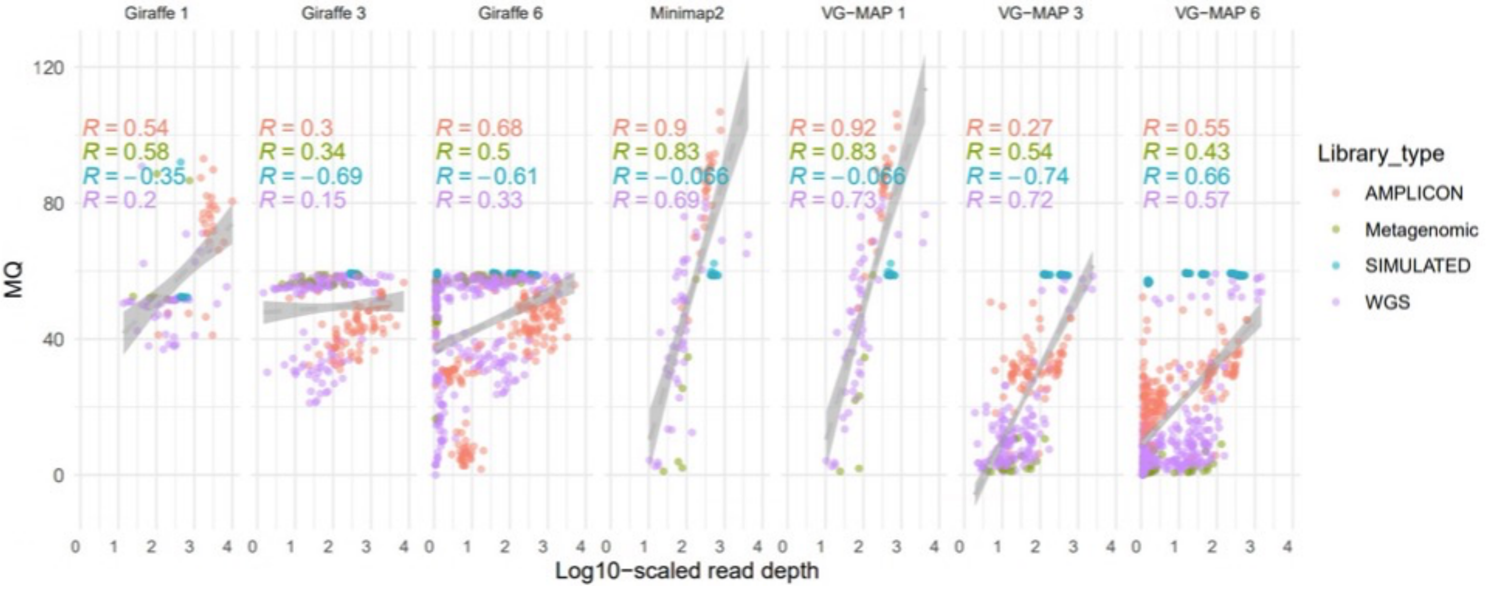


**Figure S3.1**. The effect of higher read depth (x-axis) on MQ (y-axis) for mapping approaches and library types (amplicon in red, metagenomic in green, simulated in cyan, WGS in mauve). The mappers were Giraffe, Minimap2 and VG-MAP where 1, 3 and 6 represent the 1-, 3- and 6-samples PVGs.

### 4 Supplementary Text 4: Varied mutation detection accuracy across library types, mapping tools and mutation callers

Ti/Tv ratios are positively correlated with mutation detection accuracy (1000 Genomes Project Consortium 2010, Guo et al 2012, Bainbridge et al 2011, Hodgkinson & Eyre-Walker 2010, Duchêne et al 2015). So, we evaluated the valid mutation sets for each the mapping-caller combination assuming the Minimap2 rate was the true one, which equated to needing Ti/Tv >2 based on a simple model (Figure S4.1). There were no substantial differences in Ti/Tv rates between Minimap2, VG-MAP and Giraffe mapping to one-sample references for amplicon and WGS mutations detected with BCFtools and Freebayes (Table S4.1). All had Ti/Tv > 2. This suggested no intrinsic differences in mutation quality due to the read mapper used, and either or both of BCFtools and Freebayes could be used to identify mutations (Figure S4.2). This contrasted with VG, for which one-sample PVG mapping was ineffective (Table S5).

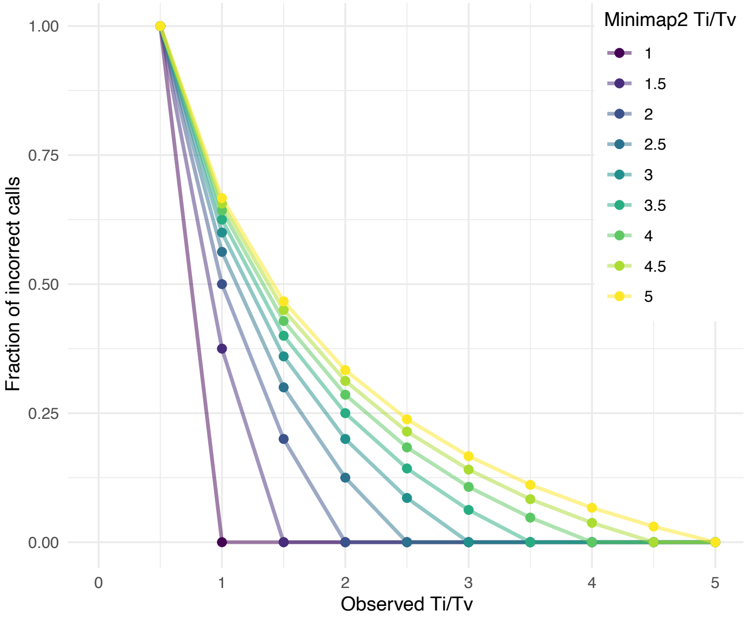


**Figure S4.1.** The effect of observed transition-transversion (Ti/Tv) rates on the estimated fraction of incorrect mutation calls given known Ti/Tv estimates from Minimap2 mapping. The Ti/Tv rate colours range from purple (Ti/Tv=1) to green (Ti/Tv=3) to yellow (Ti/Tv=5). For example, if the Ti/Tv in Minimap2 data is 3, and the observed Ti/Tv in the new dataset is 2.5, the fraction of incorrect calls is about 0.08.

Given the other results, this suggested mapping with Giraffe to the three-sample PVG was the best overall option because it had slightly higher average Ti/Tv ratios compared to the six-sample PVG (Figure S4.3). It was clear that Giraffe mapping to the three-sample PVG with BCFtools or Freebayes would be effective for WGS libraries (Table S4.1), which was also true for amplicon libraries, but for BCFtools only. Mapping to the six-sample PVG with either caller was effective for amplicon samples, but only for Freebayes for WGS libraries. The six metagenomic libraries had mean Ti/Tv ratios of 3.4 for Minimap2, 3.4 for Giraffe mapping to the one-sample PVG, and 2.3 to the three-sample PVG.

| | | Amplicon | | WGS | |
|---|---|---|---|---|---|
| Mapper | Reference | BCF | FB | BCF | FB |
| Minimap2 | Linear | 3.40 | 2.73 | 3.39 | 3.59 |
| VG-MAP | one-sample PVG | 3.40 | 3.82 | 3.46 | 3.67 |
| | three-sample PVG | 3.02 | 4.19 | 3.13 | 2.77 |
| | six-sample PVG | 3.70 | 4.22 | 1.92* | 2.20 |
| | all-sample PVG | 1.25* | 0.77* | 1.79* | 2.17 |
| Giraffe | one-sample PVG | 3.39 | 3.60 | 3.38 | 3.65 |
| | three-sample PVG | 2.05 | 1.73* | 2.72 | 2.87 |
| | six-sample PVG | 2.59 | 2.09 | 1.97* | 2.18 |
| | all-sample PVG | 0.77* | 0.91* | 1.78* | 2.07 |

**Table S4.1**. Ti/Tv rates for different library types (amplicon and WGS) based on the mapping tool (Minimap2, VG-MAP, Giraffe), PVG type (1-, 3-, 6-, all-sample or linear) and mutation caller (BCF stands for BCFtools, FB for Freebayes). *Distinctly low Ti/Tv rates.

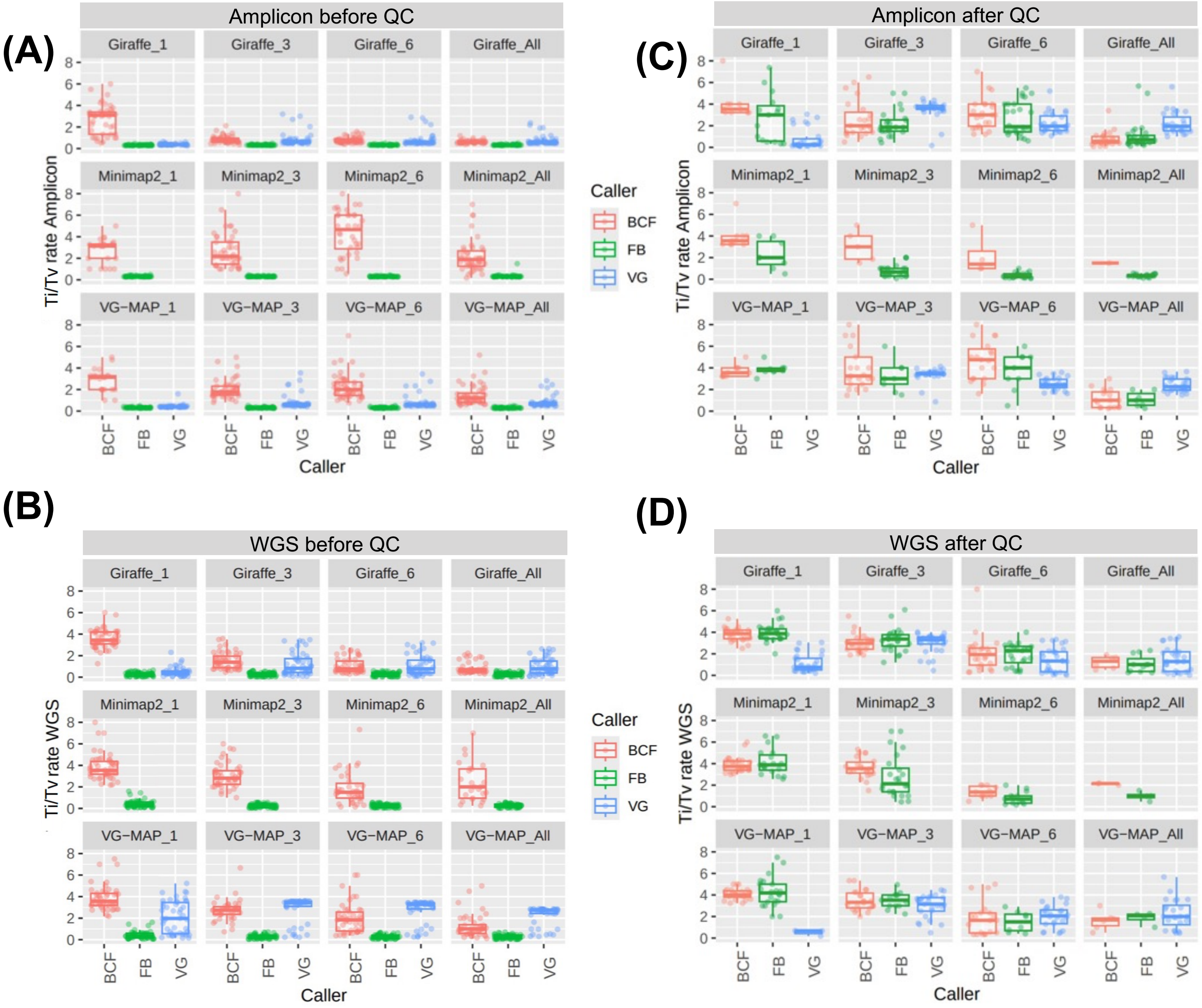


**Figure S4.2.** Ti/Tv rates for different library types, mapping tools, PVG options and mutation callers prior to and after filtering for amplicon (n=27 samples) and WGS (n=46) libraries. (A) and (C) show amplicon data before and after filtering, respectively; and (B) and (D) the same for WGS data. These showed differing effects of mapping tools, PVG options and mutations callers on Ti/Tv rates. The mapping tools are Giraffe (top of each panel), Minimap2 (middle of each) and VG-MAP (bottom of each). The PVG options were 1, 3, 6 or all samples: Minimap2 mapped to the corresponding number of samples in the reference genome file. The mutation callers were BCFtools (BCF, red), Freebayes (FB, green) and VG (blue).

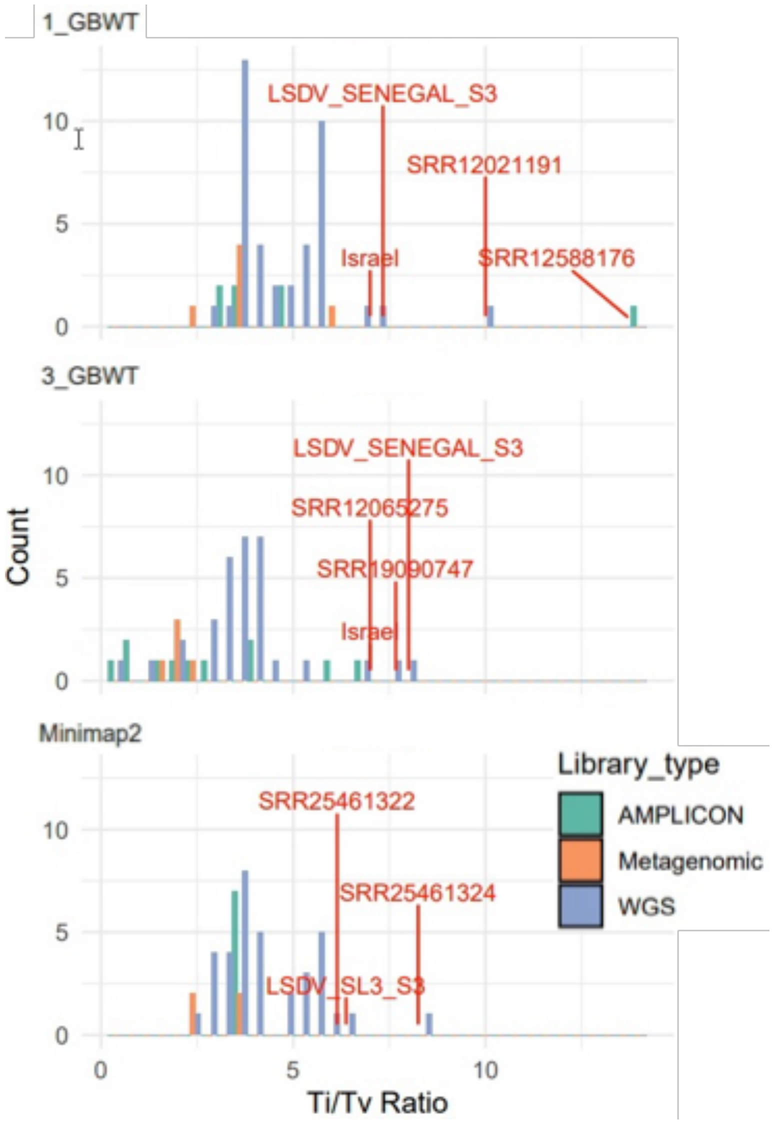


**Figure S4.3.** The Ti/Tv ratios of amplicon (green), metagenomic (orange) and WGS (blue) libraries based on valid SNPs mapping with (top) Giraffe to the 1-sample PVG, (middle) Giraffe to the 3-sample PVG, and (bottom) Minimap2 to the linear reference. Samples with Ti/Tv > 6 are highlighted in red. Six samples (Israel, Senegal_S3, SRR12021191, SRR12065275, SRR21590384 and SRR26747366) had Ti/Tv>5 in across all mapping options. 49 samples had non-zero Ti/Tv ratios in all three mapping options.

### 5 Supplementary Text 5: PVGs deliver better SNP detection in mixed samples

Mapping with Giraffe was more effective than linear mapping with Minimap2 at detecting SNPs in two mixed samples previously reported as having different levels of a Neethling-like LSDV strain, a KSGP-like LSDV strain and a Sudan-like GTPV: SRR19090747 (~99% LSDV, B-0517_PCR) and SRR19090748 (~83% LSDV, B-0517_DNA) (Vandenbussche et al 2022). Kraken2 results suggested that the SRR19090747 reads were 11% GTPV, 88% LSDV and 1% SPPV. For SRR19090748, the rates were 19%, 79% and <1%, respectively. Minimap2 mapped 73% and 86% of the reads for this pair, respectively, whereas Giraffe mapping to the one-sample PVGs mapped 99% and 93% (respectively), and mapping to the three- or six- or all-sample PVG mapped >99% of reads for all. The mixed composition of these samples was supported by the RDAF distributions of sub-consensus SNPs (Figure S5.1). BCFtools was more effective for SNP detection in SRR19090747, whereas Freebayes was better for SRR19090748. The numbers of SRR19090748 subconsensus SNPs found by Giraffe was much higher than that for Minimap2 (Figure S16). This illustrated how PVG-based methods can profile the sub-consensus SNPs of mixed samples compared to linear reference mapping, yielding information on the relative frequencies of the underlying mixed virus populations.

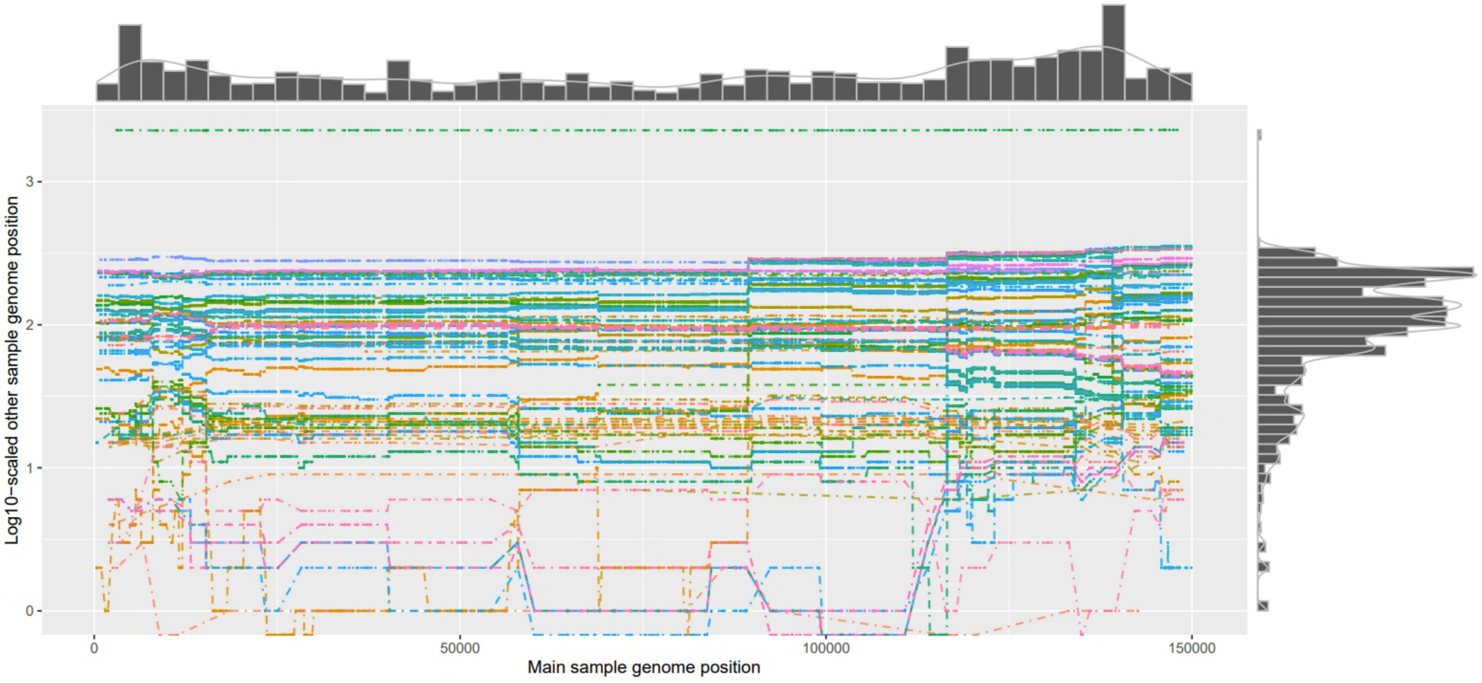


**Figure S1.** The difference in PVG coordinates across samples. Each coloured line represents one sample. The x-axis indicate the genomic position. The y-axis shows the pairwise difference in genomic positions for each sample pair (log10-scaled). MW631933 (Morocco, 14/07/2017) was the only sample with coordinates >360 bp different to the others. The histograms above and to the right of the x- and y-axes represent the frequency of each differences per locus (x-axis) and a frequency distribution of the differences (y-axis).

**Figure S2.** Allele frequency spectrum based on the all-sample PVG showing the fraction of samples (x-axis) versus the number of mutations (y-axis).

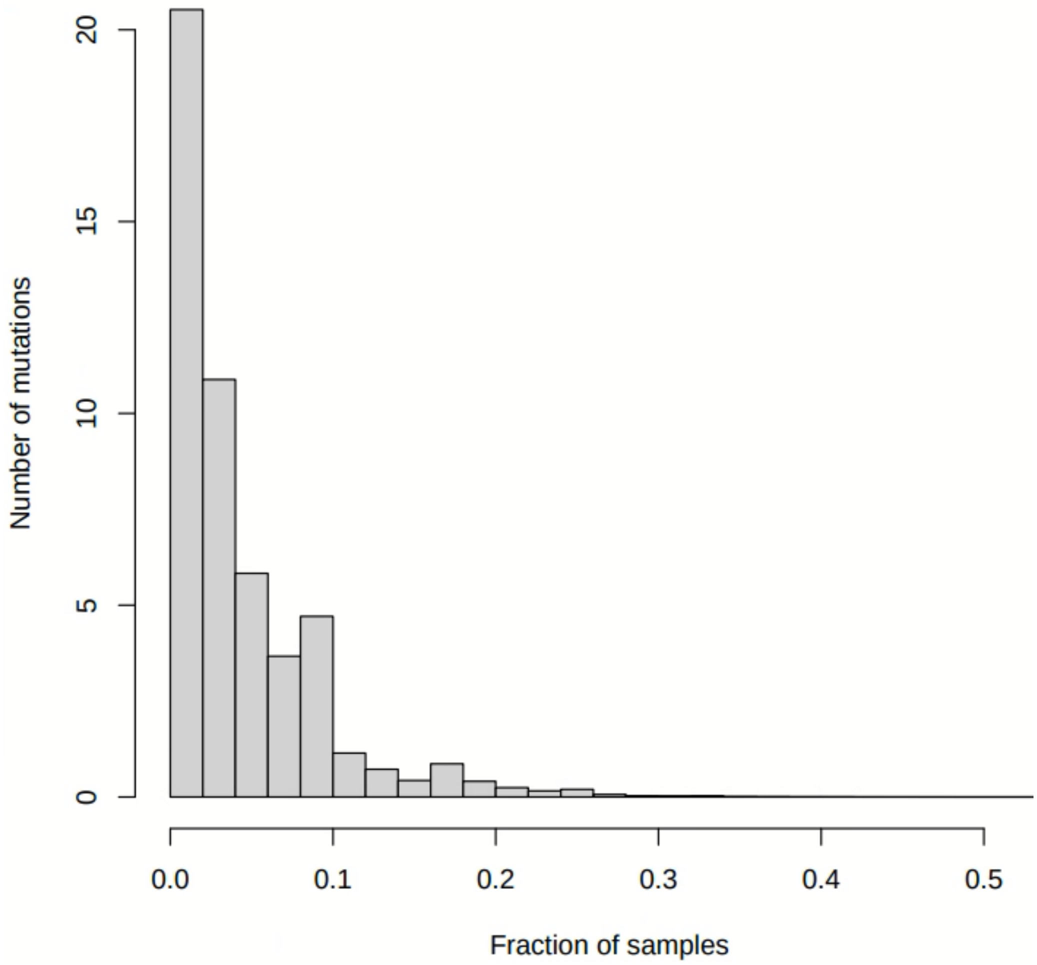

**Figure S3.** A phylogeny based on whole genome SNP data. Clade 1.1 is highlighted in navy, Clade 1.2.1 in cyan, Clade 1.2.2 in grey, Clade 1.2.3 in brown and Clade 2 in orange. Rare recombinant (R) lineages are denoted 2.1, 2.2, 2.4 and 2.6. Nodes with bootstrap support > 90 are shown. The node separating Clades 1.2.1 and 1.2.3 had 98% bootstrap support. Right: the countries are show in the first column, and the year of isolation in the second column. Both R_2.4 samples (OM530217 and OR194148, both Russia, 2019) were more like Clade 1.1, followed by R_2.1 (OL542833) and then R_2.6 (OR194148, Russia, 2019). R_2.2 (MT134042) (both Russia, 2019) was most closely related to Clade 1.2. The phylogeny was constructed with RAxML using a GTR+G4 substitution model.

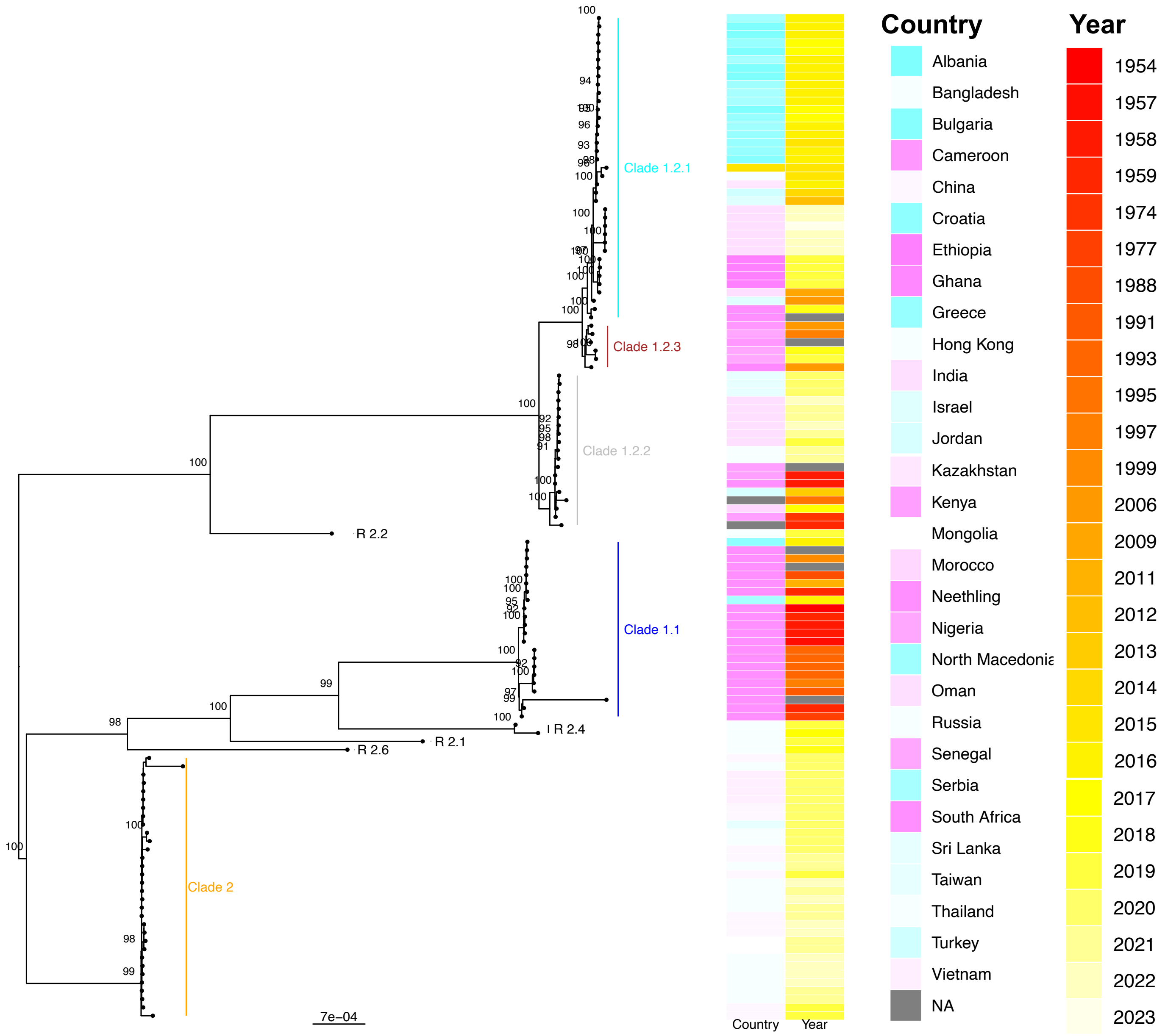

**Figure S4.** The rate of reads mapped (y-axis, %) as a function of the PVG type for amplicon libraries (n=27) mapped with Giraffe, Minimap2 or VG-MAP. The PVGs were made from either one (_1), three (_3), six (_6) or all samples (_all). The data was from (A) GAM and (B) surjected BAM files. Read mapping was best for Giraffe mapping to the one-sample PVG (red).

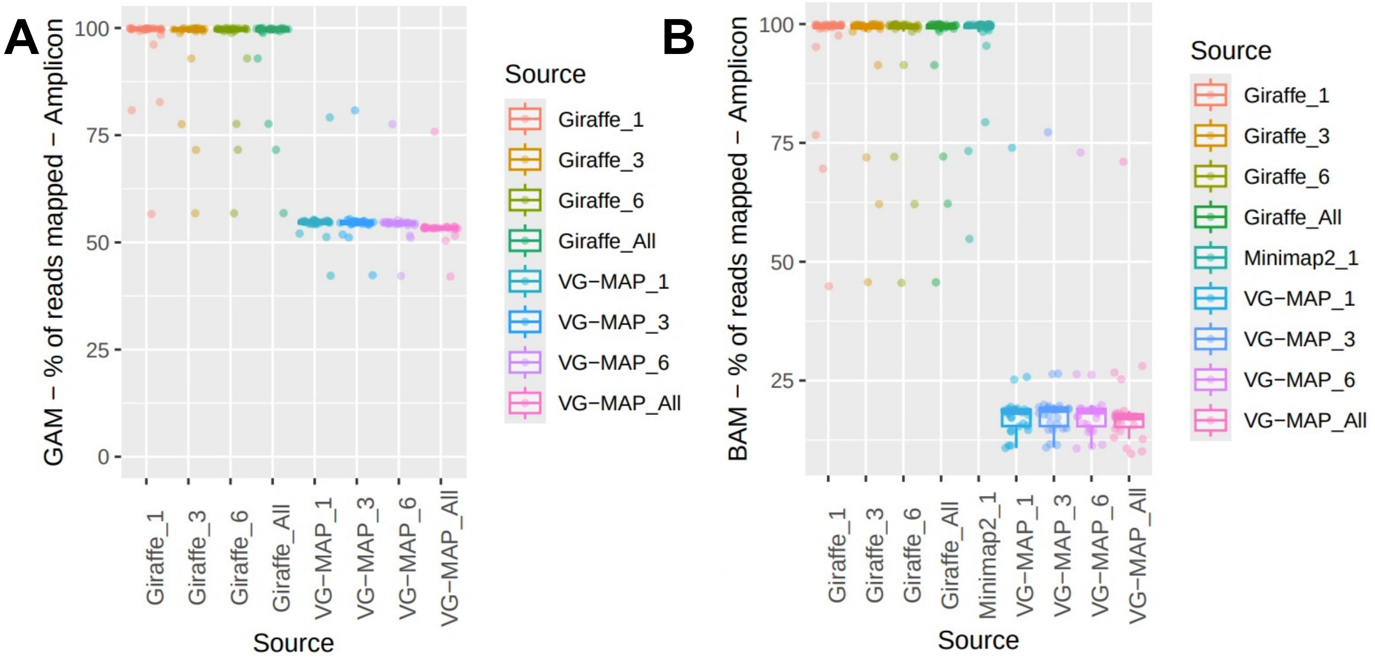

**Figure S5.** The rate of reads mapped (y-axis, %) as a function of the PVG type for Illumina WGS libraries mapped with Giraffe, Minimap2 or VG-MAP. The PVGs were made from either one (_1), three (_3), six (_6) or all samples (_all). Estimates from (A) GAM and (B) surjected BAM files.

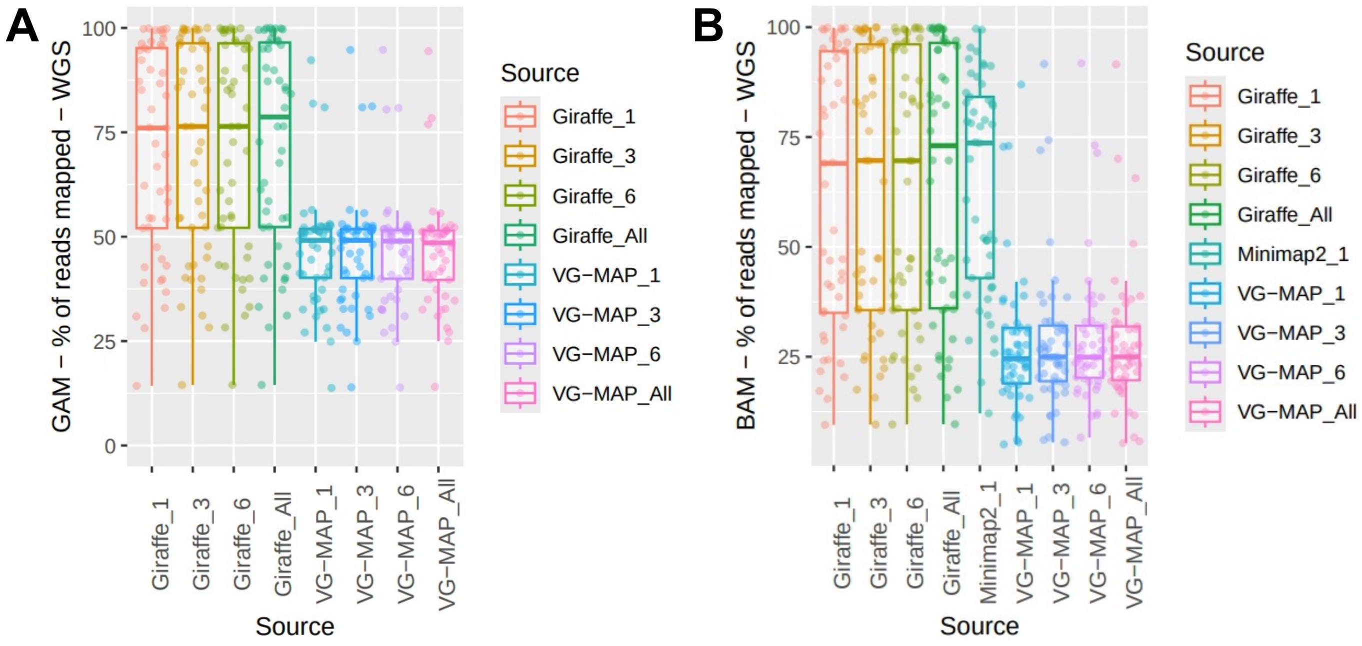

**Figure S6.** The rate of reads mapped (y-axis, %) as a function of the PVG type for Illumina metagenomic libraries (n=6) mapped with Giraffe, Minimap2 or VG-MAP. The PVGs were made from either one (_1), three (_3), six (_6) or all samples (_all). Estimates from (A) GAM and (B) surjected BAM files.

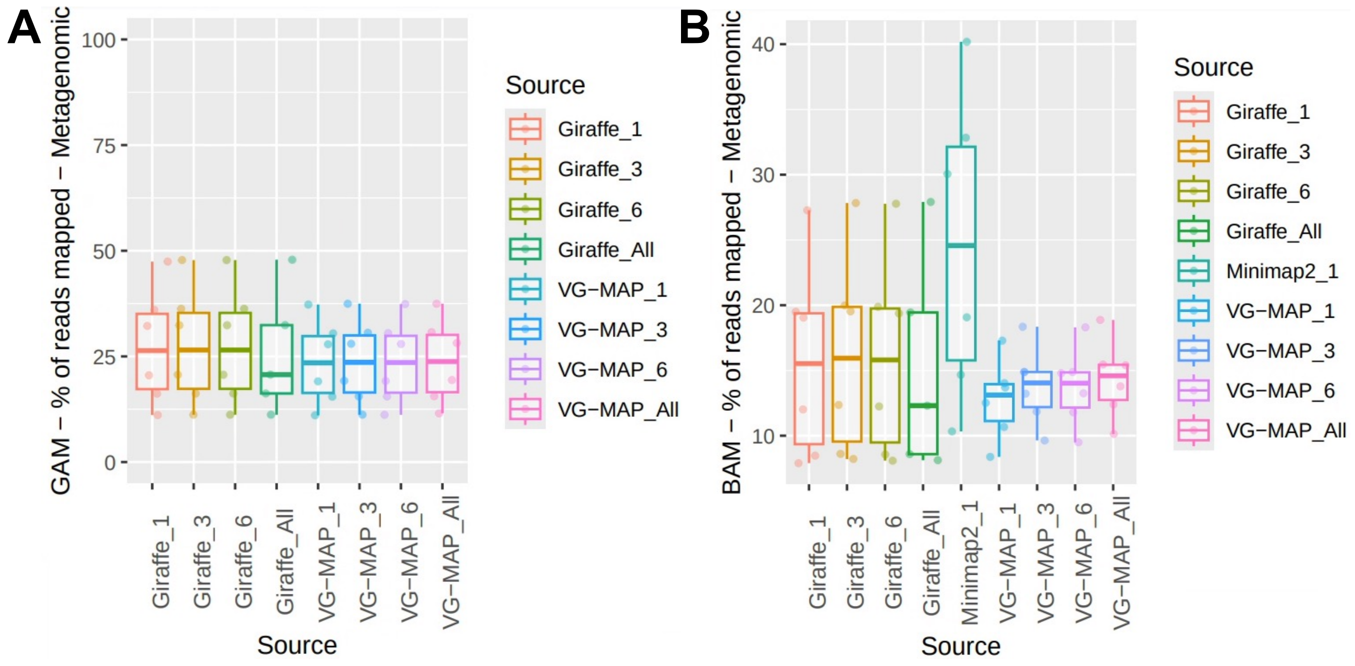

**Figure S7.** The log10-scaled numbers of screened mutations for different library types, mapping tools, PVG sizes and mutation callers. The library types (amplicon and WGS) are arrayed below the x-axis. The SNP callers on the right y-axis (BCF for BCFtools, FB for Freebayes). The panels show the numbers of mutations detected using Giraffe or Minimap2 or VG-MAP to map reads to the one-, three-, six- or all-sample PVGs. Minimap2 mapped reads to a single reference.

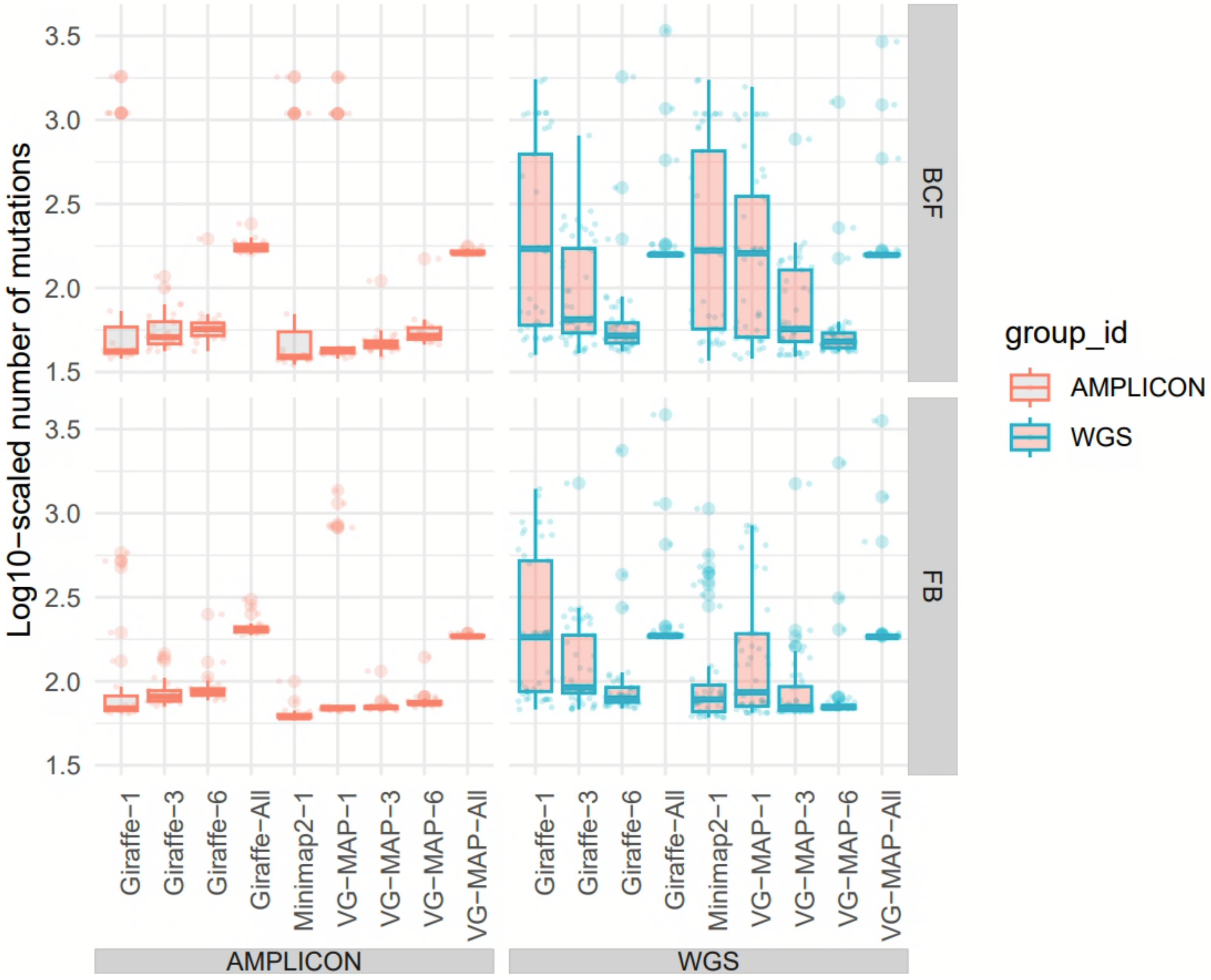

**Figure S8.** The frequencies (y-axes) of SNPs' read-depth allele frequencies (RDAF) (x-axes) for paired Illumina libraries whose reads were mapped with (1st column) Giraffe to the 1- and 3-sample PVGs (labelled GBWT_1_3), (2nd column) Giraffe to the 1- and 6-sample PVGs (GBWT_1_6), and (3rd column) Minimap2 to the linear reference (Minimap2). Red represents BCFtools (BCF), and green-blue represents Freebayes (FB).

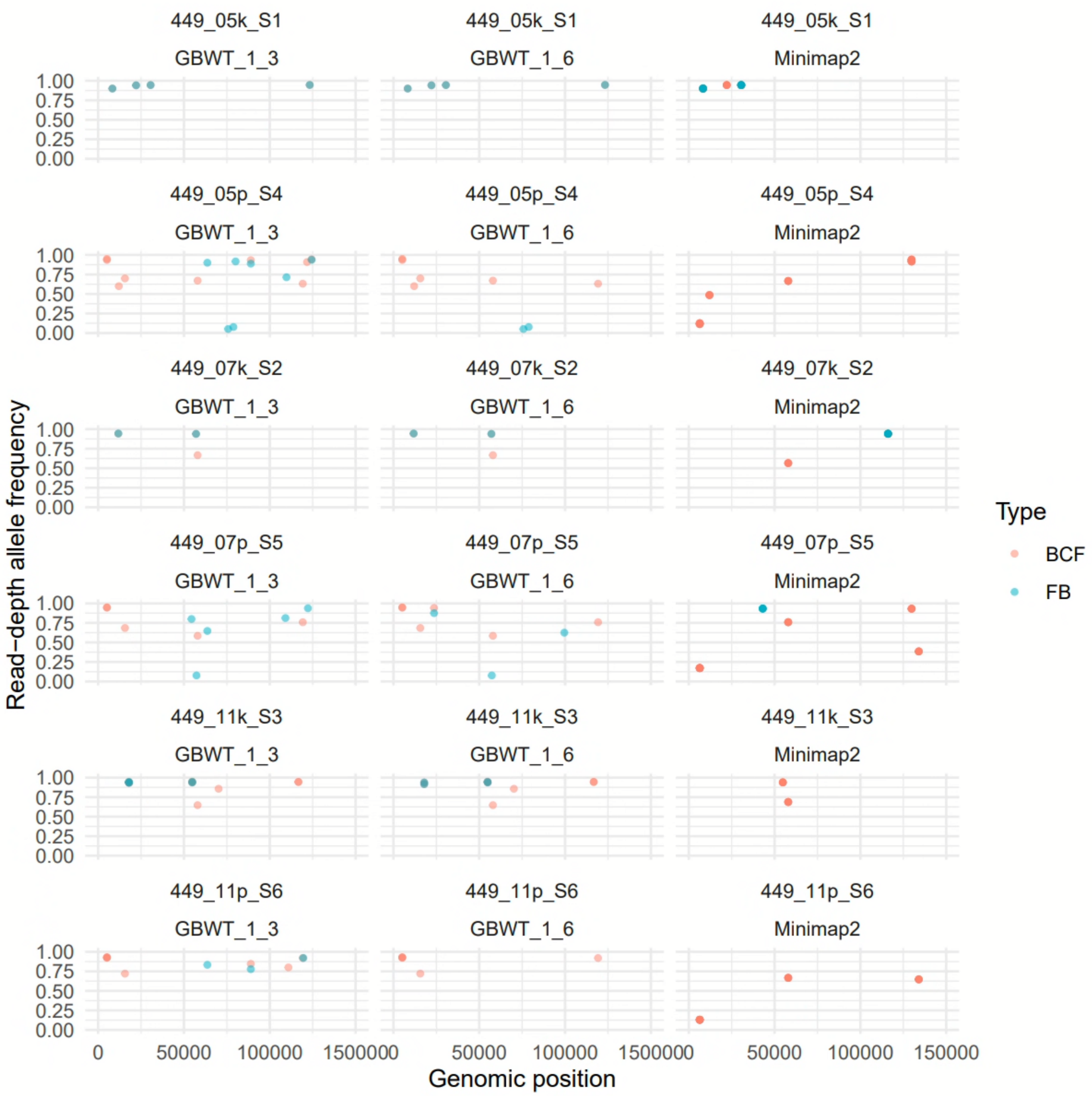

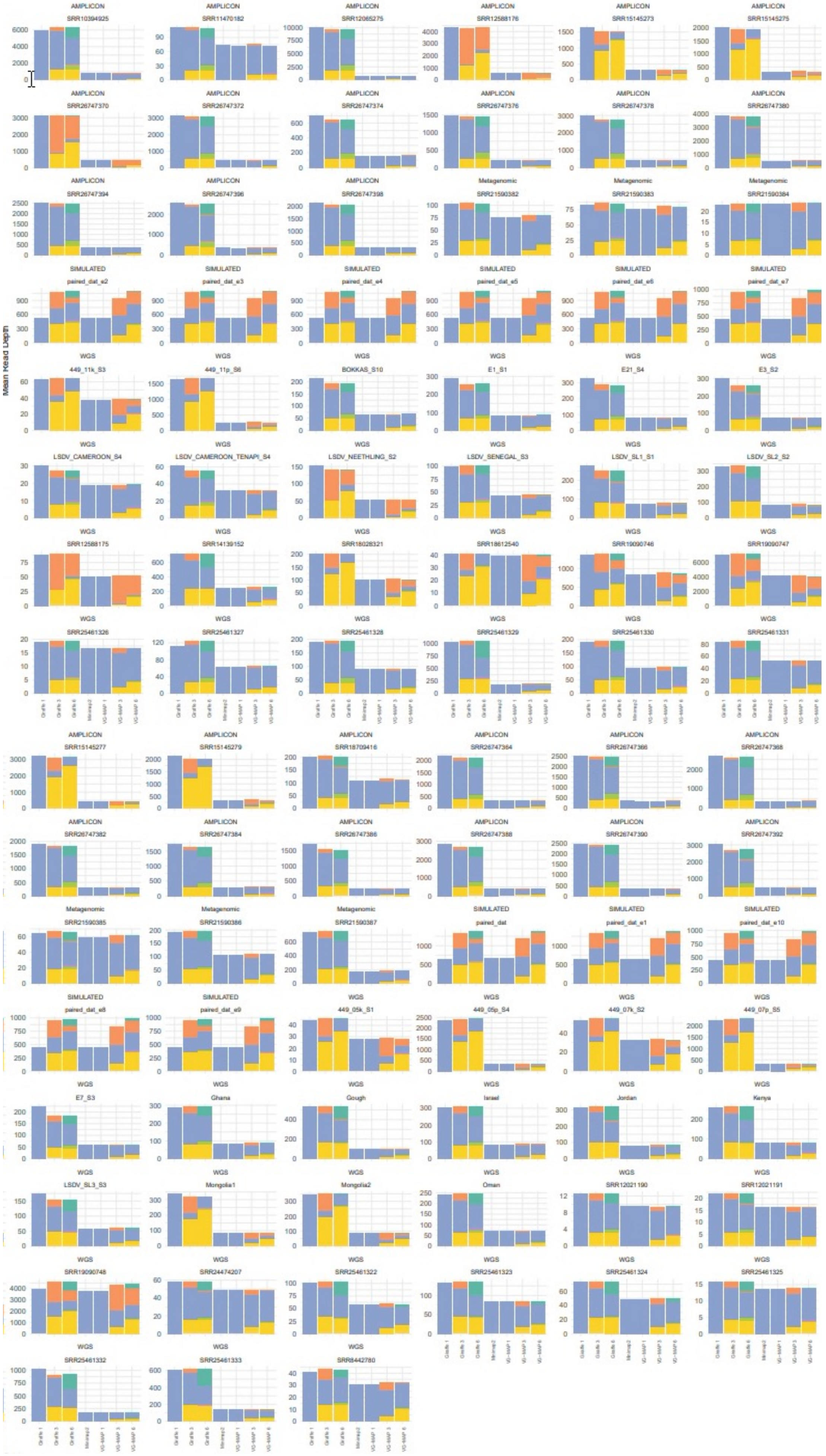

**Figure S9.** The read depth (y-axis) across different mapping approaches and reference samples (KX683219 in aquamarine, KX764645 in orange, KX894508 in blue, MW732649 in mauve, OK318001 in moss and OQ511520 in mustard). The mappers were Giraffe, Minimap2 and VG-MAP where 1, 3 and 6 represent the 1-, 3- and 6-sample PVGs.

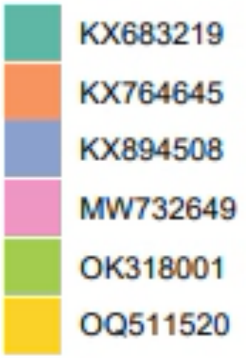

**Figure S10.** Visualisations of the LSDV PVG at different scales. (A) The PVG topology at 5.2-6.5 Kb where the path bifurcations represent alternate allelic combinations visualised using Bandage-NG's force-directed layout where the contig colours are randomly assigned for visual clarity. LD008 (encoding a putative soluble interferon gamma receptor) was at 4,842-5,669 bp; LD009 (encoding a putative alpha amanitin-sensitive protein) was at 5,702-6,394 bp. (B) SequenceTubeMap image of the 121-sample PVG at KX894508:5,831-5,897 bp (spanning LD008 and LD009) showing seven SNPs (0.105 SNPs/Kb) with two major haplotypes with 56 (top) and 65 (bottom) samples.

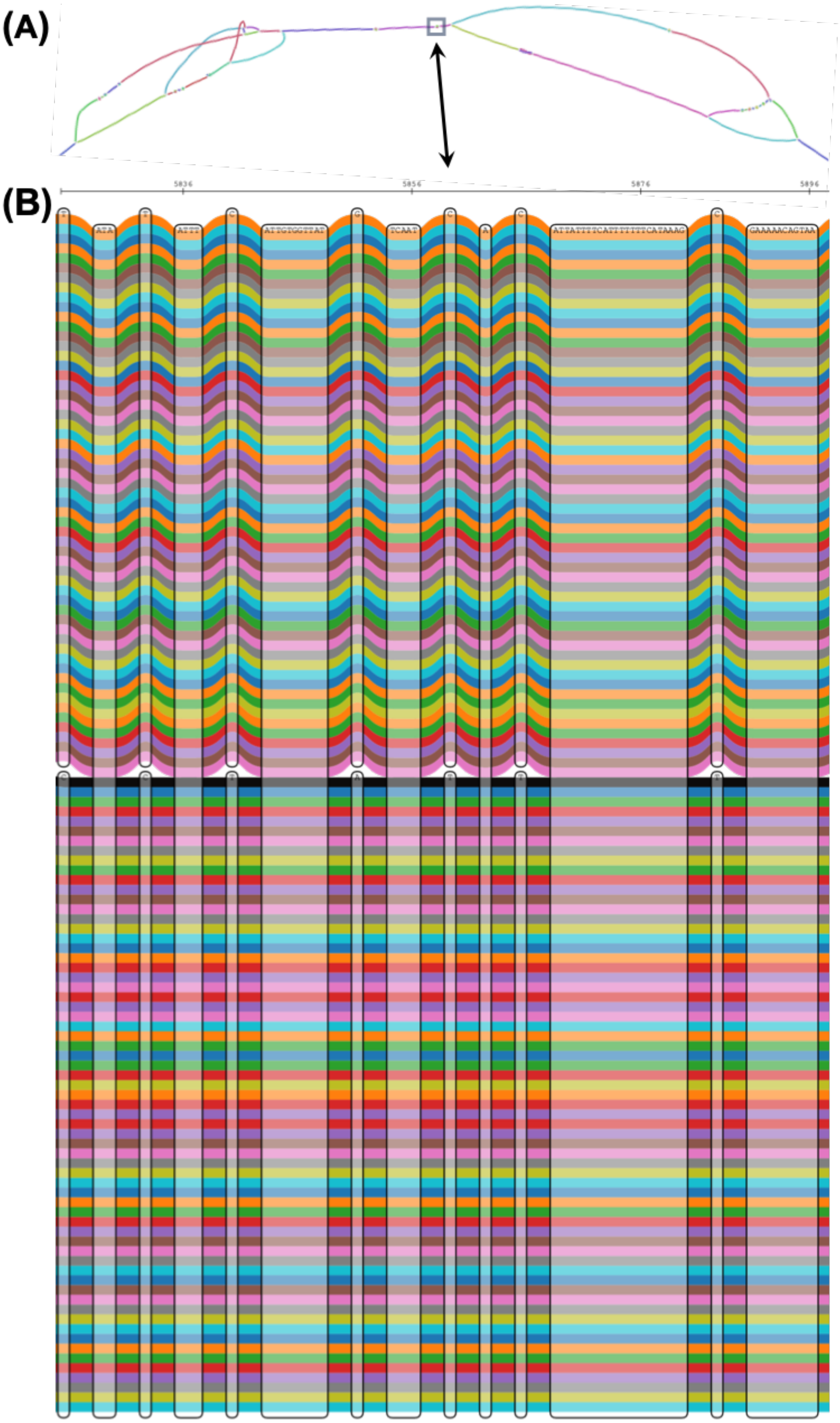

**Figure S11.** Visualisations of the LSDV PVG at different scales. (A) VG visualisation of 8,055-8,231 bp with nine mutations (0.051 SNPs/Kb). LD011 (encoding a CC chemokine receptor-like protein) was at 6,980-8,113 bp. (B) SequenceTubeMap image of 449_07p_S5's reads (red and blue) mapped to the three-sample PVG at 8,062-8,131 bp (at LD011) showing three SNPs where KX894508 (Clade 1.2) is in light grey, OQ511520 (Clade 2) is in grey, and KX764645 (Clade 1.1) is coloured black.

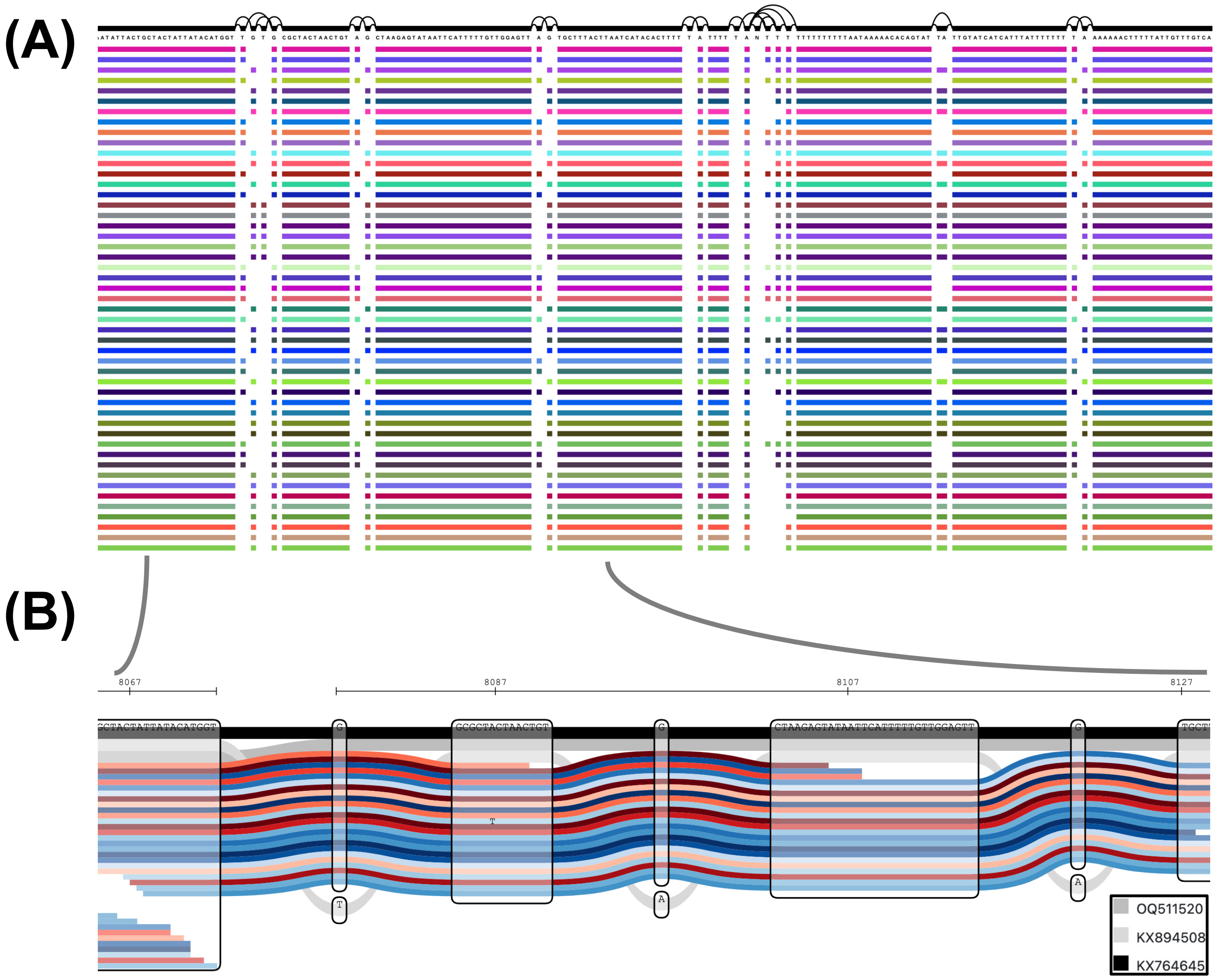

**Figure S12.** Visualisations of the LSDV PVG at different scales. (A) ODGI image of 136,200-140,400 bp. LD144 (encoding a kelch-like protein) was at 135,541-137,208 bp. LD145 (encoding an ankyrin repeat protein) was at 137,232-139,136 bp. LD146 (encoding a phospholipase D-like protein) was at 139,265-140,506 bp. (B) SequenceTubeMap image of 449_07p_S5's reads (red and blue) mapped to the three-sample PVG at 140,272-297 bp showing five SNPs where KX894508 (Clade 1.2) is in light grey, OQ511520 (Clade 2) is in grey, and KX764645 (Clade 1.1) is coloured black.

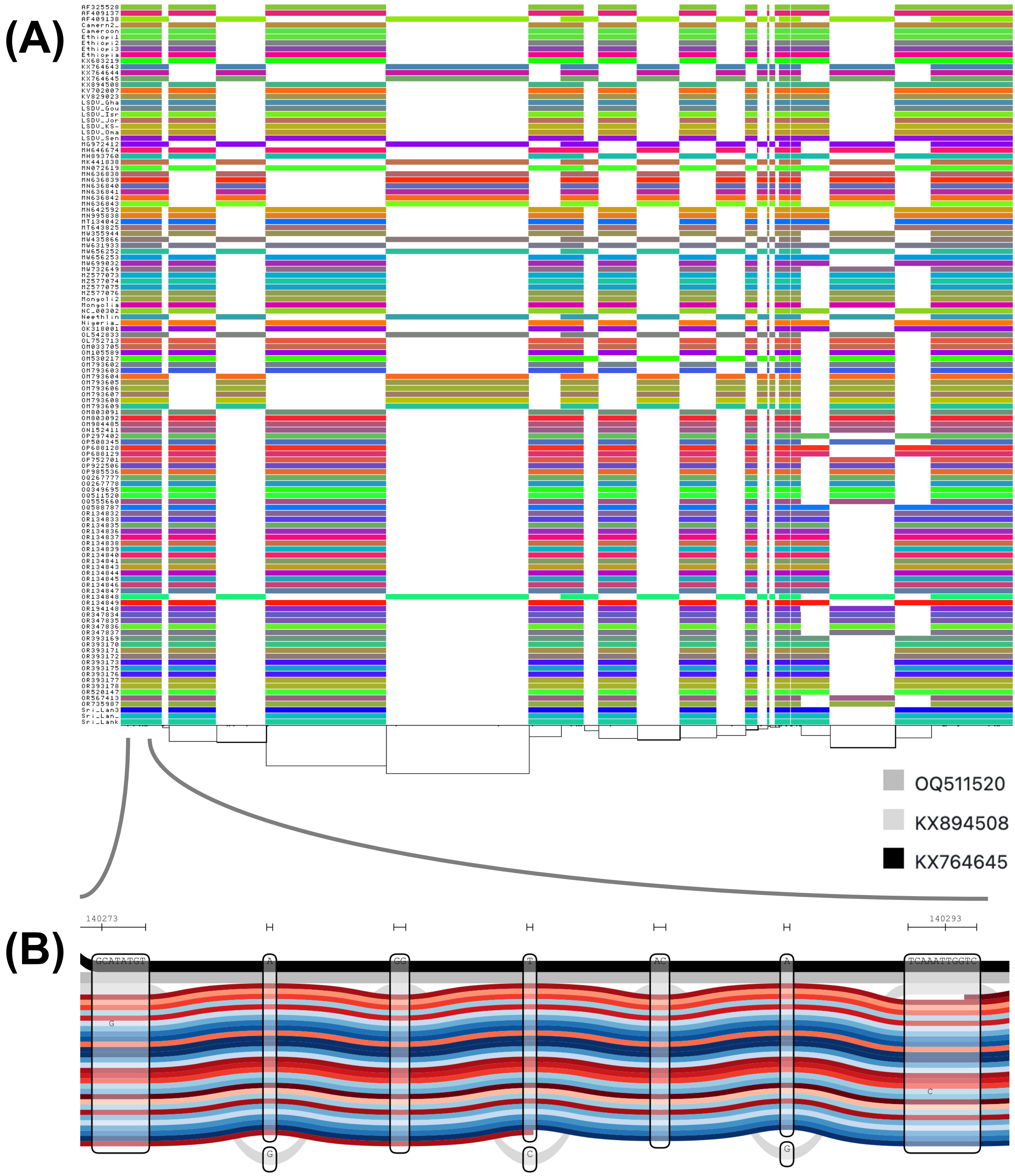

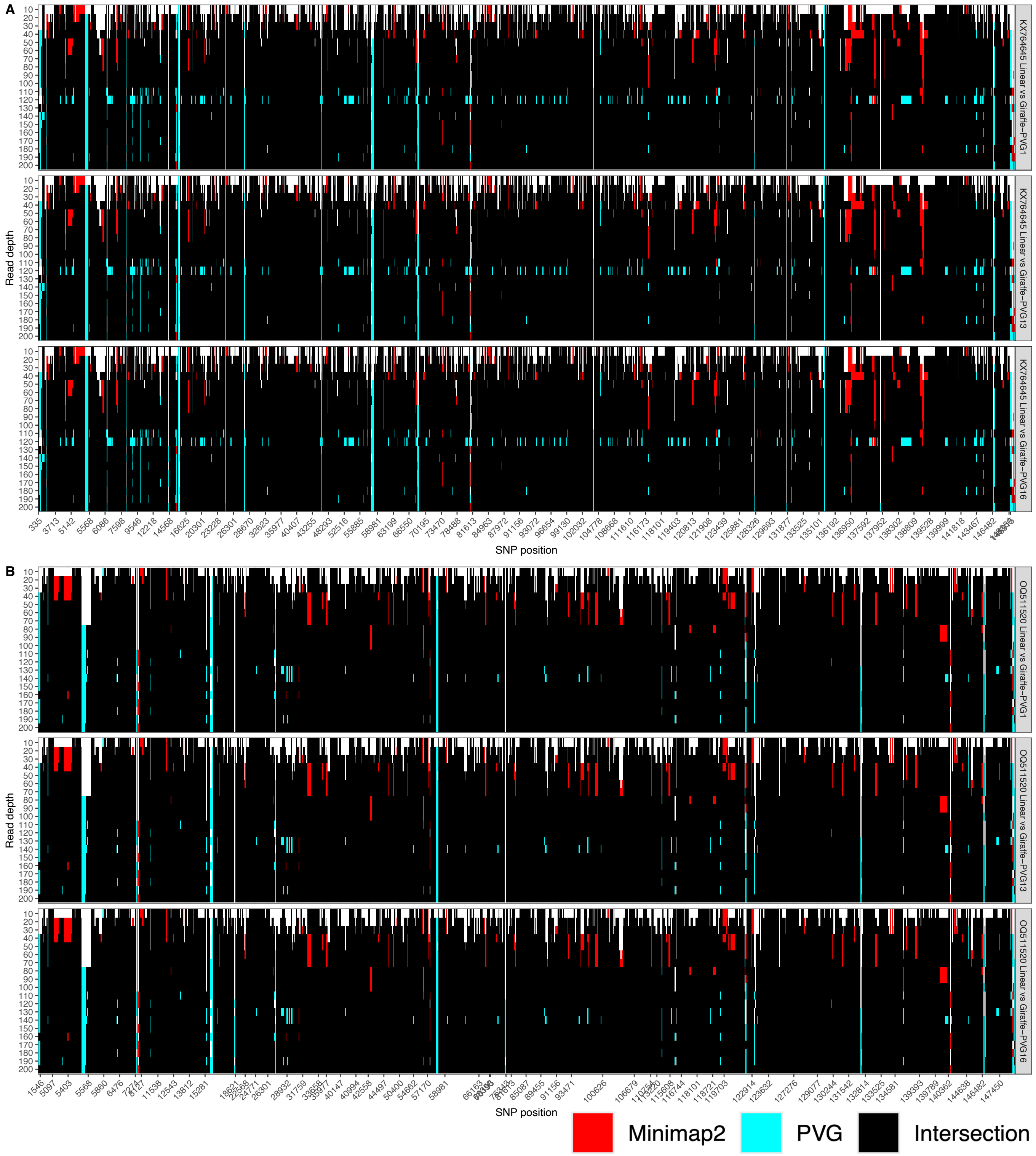


**Figure S13**. Rates of genome-wide SNPs detection as a function of read depth (y-axis) using simulated reads based on (A) KX764645 and (B) OQ511520. In (A) and (B), the panels show the SNPs detected using Giraffe mapping to the: (top) one-sample PVG, (middle) combined one- and three-sample PVGs and (bottom) combined one- and six-sample PVGs. SNPs with a red colour indicates SNPs detected by Minimap2 alone. Cyan indicates those detected by PVG-based mapping only. Black indicates those found by both Minimap2 and PVG mapping.

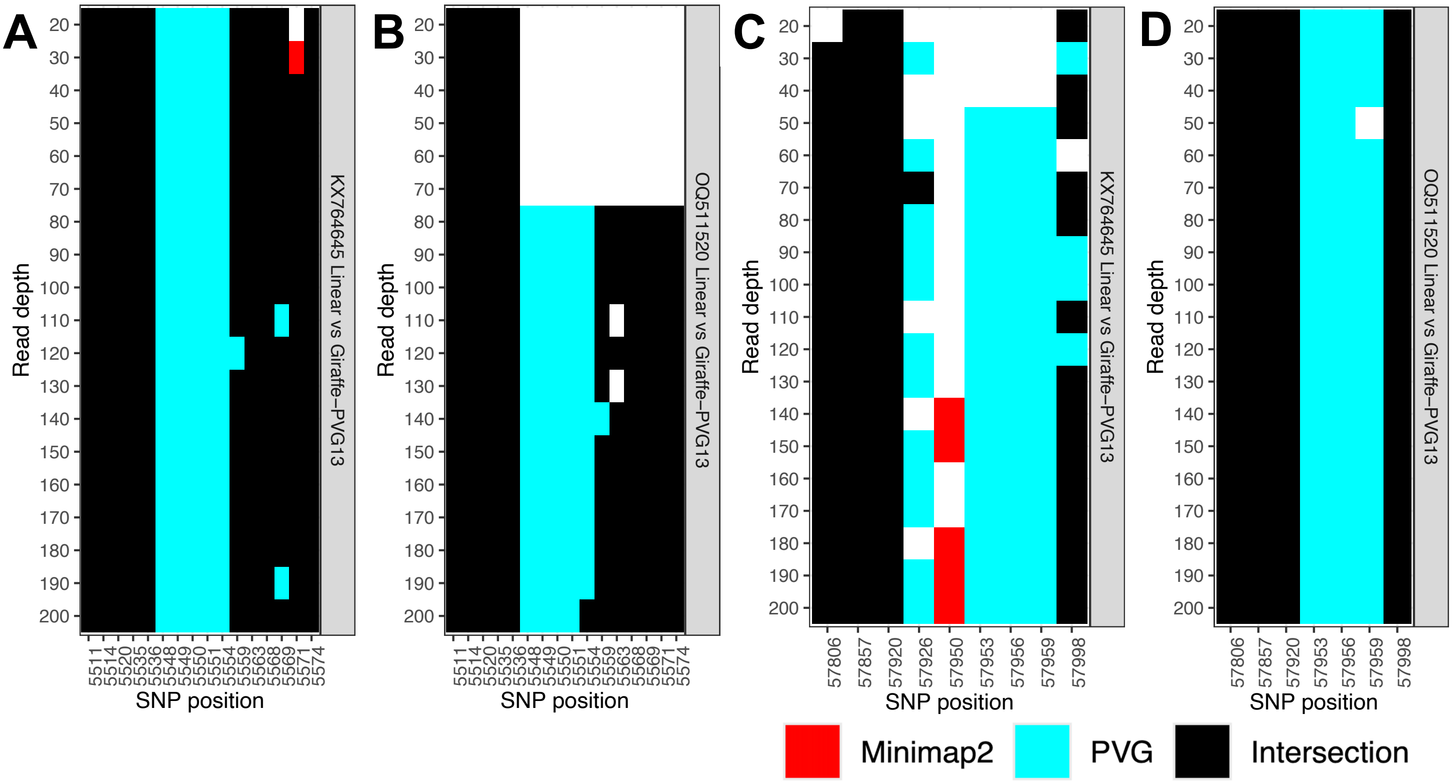


**Figure S14.** Nonsynonymous SNP clusters at regions (A/B) 5,500-5,600 bp and (C/D) 57,800-58,000 bp as a function of read depth. The SNP data comes from simulated reads based on (A/C) KX764645 and (B/D) OQ511520. SNPs with a red colour indicates SNPs detected by Minimap2 alone. Cyan indicates those detected by PVG-based mapping only. Black indicates those found by both Minimap2 and PVG mapping. The SNPs detected by PVG-based mapping at (A) 5,548, 5,549, 5,551, 5,554, (B) 57,953, 57,965 and 57,959 bp all caused nonsynonymous effects.

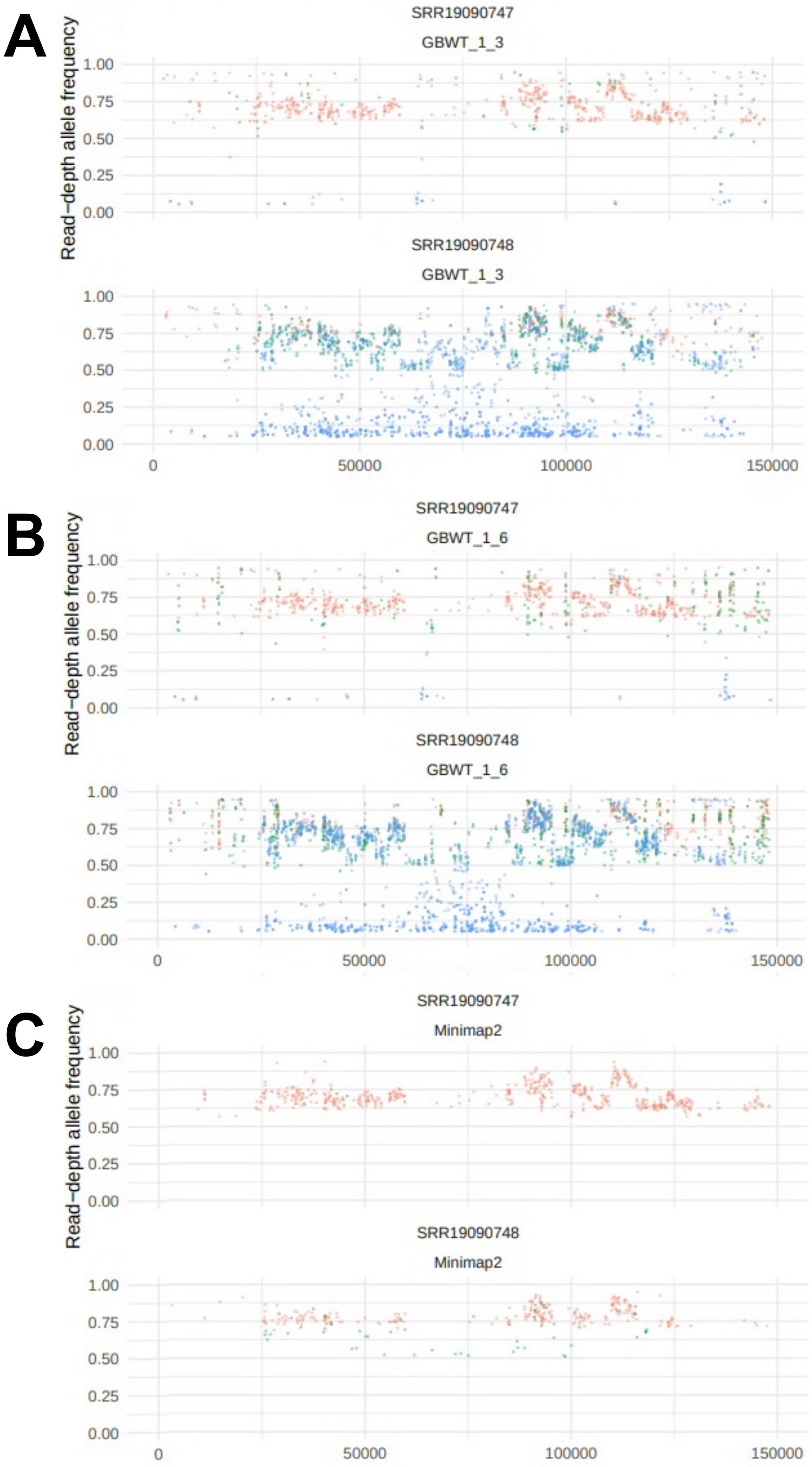


**Figure S15.** The read-depth allele frequencies (RDAF) of SNPs (y-axes) across the genome (x-axes) for the two Lumpivax mixed samples whose reads were mapped with (A) Giraffe to the 1- and 3-sample PVGs (labelled GBWT_1_3), (B) Giraffe to the 1- and 6-sample PVGs (GBWT_1_6), and (C) Minimap2 to the linear reference (Minimap2). The points are coloured red for those detected by BCFtools (BCF), green for Freebayes (FB) and blue for VG (VG). The samples were SRR19090747 (B-0517_PCR) and SRR19090748 (B-0517_DNA).

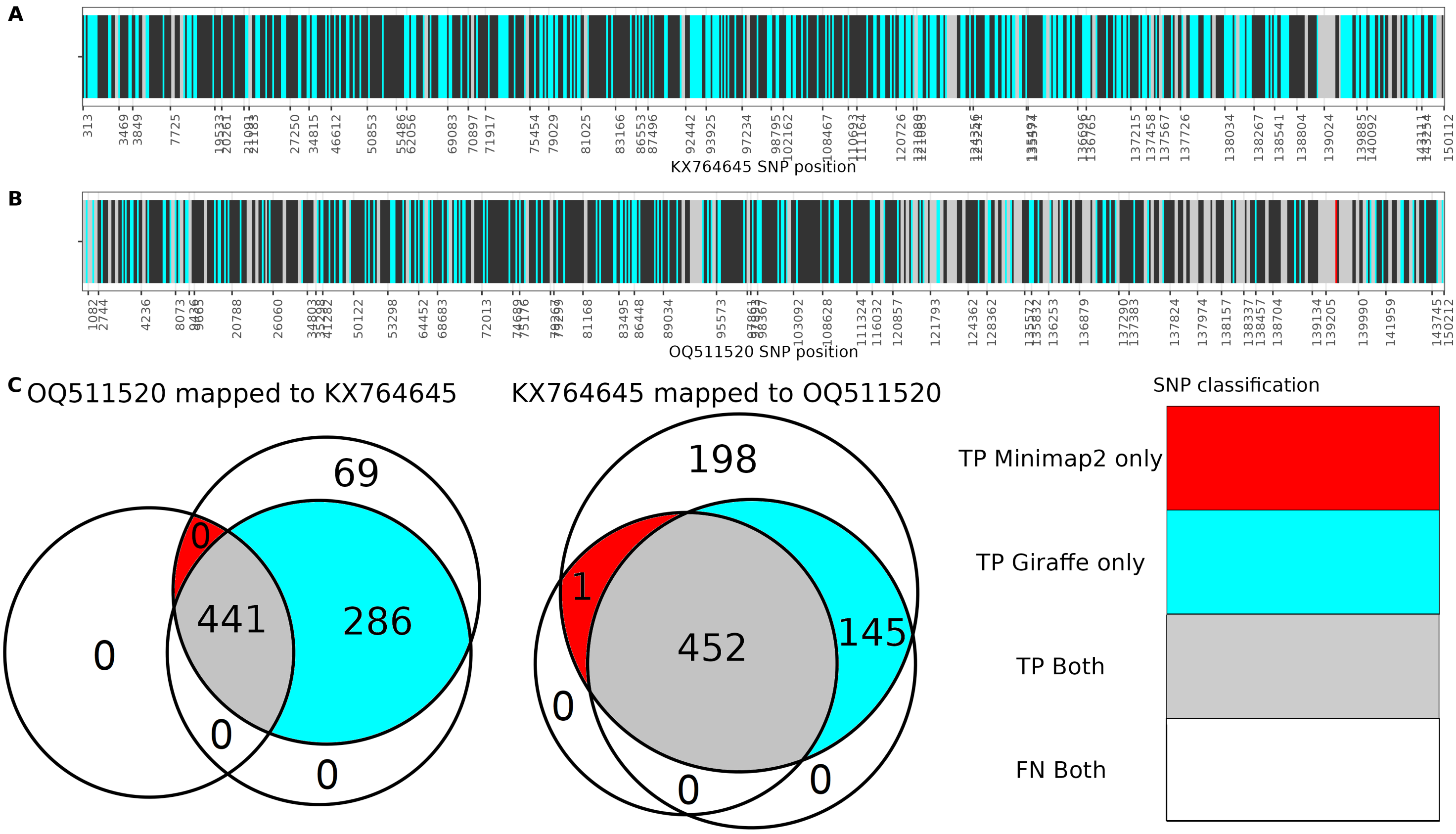


**Figure S16.** Genome-wide SNP detection for simulated reads from (A) OQ511520 mapped to reference KX764645 and from (B) KX764645 mapped to reference OQ511520. SNPs detected by both mapping methods (Minimap2 and Giraffe) are in black, those detected by Giraffe only are in cyan, those detected by Minimap2 only are in red, and those missed by both Giraffe and Minimaps are in grey. (C) Venn diagrams of the SNP numbers unique to each set: Minimap2, Giraffe, and the true SNP sets, where the grey area are true positive SNPs detected by both Giraffe and Minimap2, the red area is for SNPs detected by Minimap2 only, the cyan area is for the SNPs detected by Giraffe only, the white non-zero area are the SNPs missed by both Minimap2 and the other white areas reflect false positive SNPs (none were found). The OQ511520 mapped to reference KX764645 is on the left, and KX764645 mapped to reference OQ511520 is on the right.

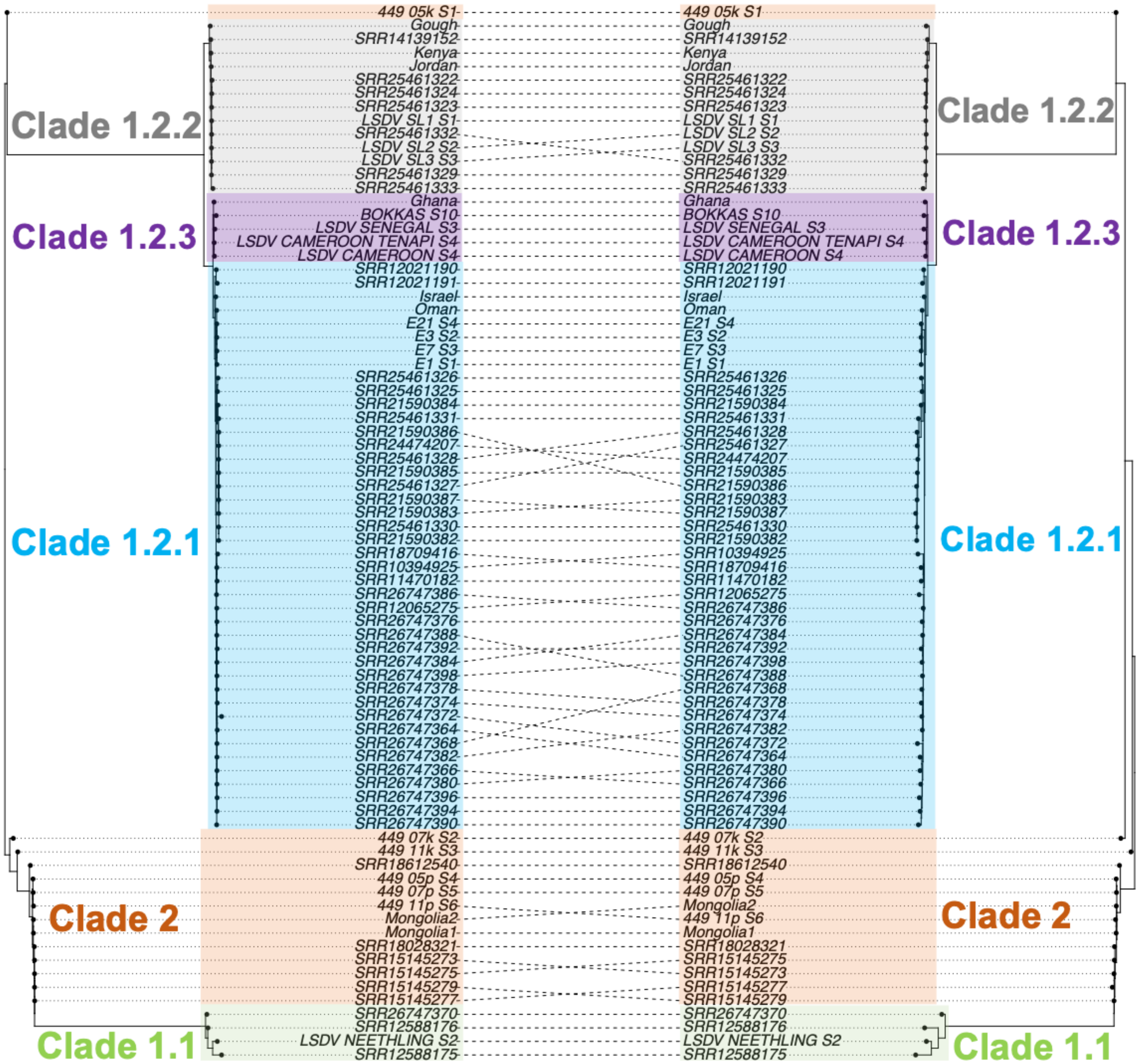


**Figure S17.** A co-phylogeny of genome-wide SNPs inferred from (left) linear reference mapping and (right) PVG-based mapping. The clades are shown by shaded areas: Clade 1.1 in pale green, subclade 1.2.1 in pale cyan, subclade 1.2.2. is in grey, subclade 1.2.3 is in purple, and Clade 2 in pale orange.
